# Cytosine base editing workflow for quality-controlled multiplex-knockout hiPSC lines

**DOI:** 10.64898/2026.06.02.729559

**Authors:** Vanessa Kirschner, Julia Herzberg, Christin Richter, Sayari Bhunia, Rebecca Kistler, Roger Ottenheijm, Timon Seeger, Marc Freichel, Alex Cornean

## Abstract

Dissecting polygenic disease mechanisms requires human cell models that harbour multiple targeted genetic modifications in a defined background. However, generating and rigorously validating such models remains difficult. We developed a cytosine base editing workflow to generate multiplex-knockout (KO) human induced pluripotent stem cell (hiPSC) lines. First, we assessed six cytosine base editor (CBE) variants and selected evoBE4max. We then combined sgRNA-guided introduction of premature termination codons and splice-site mutations with fluorescence-based enrichment. This yielded a median on-target C-to-T editing efficiency of 77.5% (range, 27.0-86.5%) across six loci. We generated single-, double-, and triple-KO hiPSC lines for endolysosomal Ca²⁺ signalling components (*OCaR2*, *TPC1*, *TPC2*) and confirmed loss-of-function at transcript and protein levels. We performed extensive quality control, including pluripotency assessment, SNP-array karyotyping, and whole-genome sequencing, which indicated minimal guide-directed off-target editing. We further extended multiplex editing to *ORAI* Ca²⁺ channel paralogs. This framework supports scalable production of quality-controlled multiplex-KO hiPSC lines.

## INTRODUCTION

Human induced pluripotent stem cells (hiPSCs) provide a powerful platform for studying human development and disease within a genetically defined, ethically sustainable, and renewable cellular system. They have widespread uses in disease modelling, drug screening, and regenerative medicine, including advanced systems like organoids and microphysiological platforms (Cerneckis et al., 2024; Ingber, 2022). A major benefit of hiPSC-based models is the ability to create isogenic lines through CRISPR-Cas9-mediated gene editing, either by gene knockout (KO) or by introducing defined genetic variants. This allows detailed analysis of genotype-phenotype relationships (Hockemeyer and Jaenisch, 2016; Sayed et al., 2016; Soldner and Jaenisch, 2018; Soldner et al., 2011). This is particularly relevant for polygenic disorders, where multiple genetic risk loci interact, and single-gene perturbations may not capture disease-relevant biology (Cerrone et al., 2019; Visscher et al., 2021). However, hiPSCs are inherently difficult to edit and expand clonally, making the efficient production of multiplex-edited lines particularly challenging (Hockemeyer and Jaenisch, 2016; Ihry et al., 2018; Standage-Beier et al., 2019; Sürün et al., 2020). Many common CRISPR methods involve inducing DNA double-strand breaks (DSBs), but these can reduce editing efficiency in hiPSCs because of p53-related toxicity and selection bias during clonal derivation (Ihry *et al*., 2018). These factors make it difficult to scale up the generation of multiplex-edited hiPSC clones for further analysis or differentiation.

Nickase-based base editors offer an alternative approach to gene disruption that does not depend on targeted DSBs. Cytosine base editors (CBEs) enable programmable C-to-T conversions in genomic DNA without requiring DSBs or donor DNA templates. These edits can be used to fix or introduce genetic variants (Komor et al., 2016). Furthermore, C-to-T transition mutations are ideal for creating predicted loss-of-function (LoF) alleles by introducing premature termination codons (pmSTOP/PTC strategies) (Billon et al., 2017; Kuscu et al., 2017), or by disrupting splice donor (SD)/acceptor (SA) motifs (Gapinske et al., 2018; Yuan et al., 2018), which can lead to measurable transcript-level effects such as transcript degradation, exon skipping or intron retention (Kluesner et al., 2021). These features make base editing particularly suitable for multiplex KO engineering (Brookhouser et al., 2020; Webber et al., 2019), where efficient selection of edited clones is essential.

A crucial element for scalable base editing is enriching edited cells prior to clonal isolation. The transient reporter for editing enrichment (TREE) connects base editor activity to a fluorescence switch, enabling the selection of edited populations by fluorescence-activated cell sorting (FACS) and enhancing the recovery of edited cells (Standage-Beier *et al*., 2019). TREE-based workflows have been adapted for hiPSCs, supporting efficient creation of isogenic lines and multiplex editing at independent loci. Brookhouser et al. developed BIG-TREE, using an episomal reporter with AncBE4max for effective clonal base editing in hiPSCs, including PTC-based gene KO, supported by representative karyotyping and pluripotency/trilineage marker analyses for these single-KOs, and importantly, demonstrated multiplex editing at independent loci (Brookhouser *et al*., 2020). More recently, Zhang et al. expanded this concept into a semi-automated iSTOP pipeline, focused primarily on PTC-mediated LoF across multiple genes and lines, emphasising the streamlined derivation of single-gene LoF alleles with systematic quality control (QC) (Zhang et al., 2024b). Despite these advances, a comprehensive end-to-end workflow for multiplex (two or more loci) LoF editing in hiPSCs remains less defined. Such a workflow should integrate modern editor and guide RNA selection, include structured assessment of pluripotency and genome integrity, demonstrate concurrent editing within the same clone, offer practical guidance on guide RNA design and selection, and realistic clone yield expectations.

We address this gap by establishing an integrated multiplex base editing workflow for clonal hiPSC engineering. We benchmarked recent CBE variants at HEK293T test loci with amplicon sequencing, then combined TREE-based enrichment and systematic sgRNA selection to create single-, double-, and triple-KO hiPSC lines using a mixture of PTC and splice-site strategies. These methods targeted three genes involved in a mechanosensitive Ca²⁺ release pathway from acidic stores: the organellar Ca²⁺ regulator *TMEM63B*/*OCaR2* (encoding OCaR2) and the two-pore channels *TPCN1* and *TPCN2* (encoding TPC1 and TPC2). Dysregulation of these genes has been associated with impairments of numerous physiological functions, including thirst, hearing and surfactant release (Du et al., 2020; Freichel et al., 2024; Yang et al., 2024; Zou et al., 2025), as well as diseases such as viral infections, cardiac arrhythmias, fatty liver disease and other Ca²⁺-related diseases (Klingl et al., 2025). However, their roles in human cellular models are still not fully understood. We evaluated edited clones using multi-layer QC, including pluripotency markers, differentiation potential, and genome integrity screening (SNP-array karyotyping and, for some lines, whole-genome sequencing), along with genotype confirmation and transcript/protein analyses consistent with the predicted LoF. Lastly, we demonstrate the approach’s portability by extending multiplex editing to ORAI Ca²⁺ channel paralogs (*ORAI1-3*) (Prakriya and Lewis, 2015). Together, this workflow enables the scalable generation of multiplex-edited hiPSC lines for mechanistic studies in human cell types.

## RESULTS

### Benchmarking cytosine base editors in human cells

To select a suitable CBE for multiplex gene disruption in hiPSCs, we prioritised broad editing windows and reliable performance across dinucleotide contexts. We compared six CBE variants - evoBE4max (Thuronyi et al., 2019), AncBE4max (Koblan et al., 2018), BE4-Gam (Komor et al., 2017), hyBE4max (Zhang et al., 2020), TadCBEd (Neugebauer et al., 2023) and CBE-T1.46 (Lam et al., 2023) - across four target loci (Chen et al., 2023a; Chen et al., 2023b) in HEK293T cells. We characterised C-to-T conversion rates for cytosines along the protospacer (Figure 1A), peak base editing efficiency, defined as the highest measured C-to-T editing within the protospacer regardless of the presence or absence of bystander editing (Figures 1B and 1D), and indel frequency (Figures 1C and 1E) by amplicon deep sequencing.

**Figure 1.**
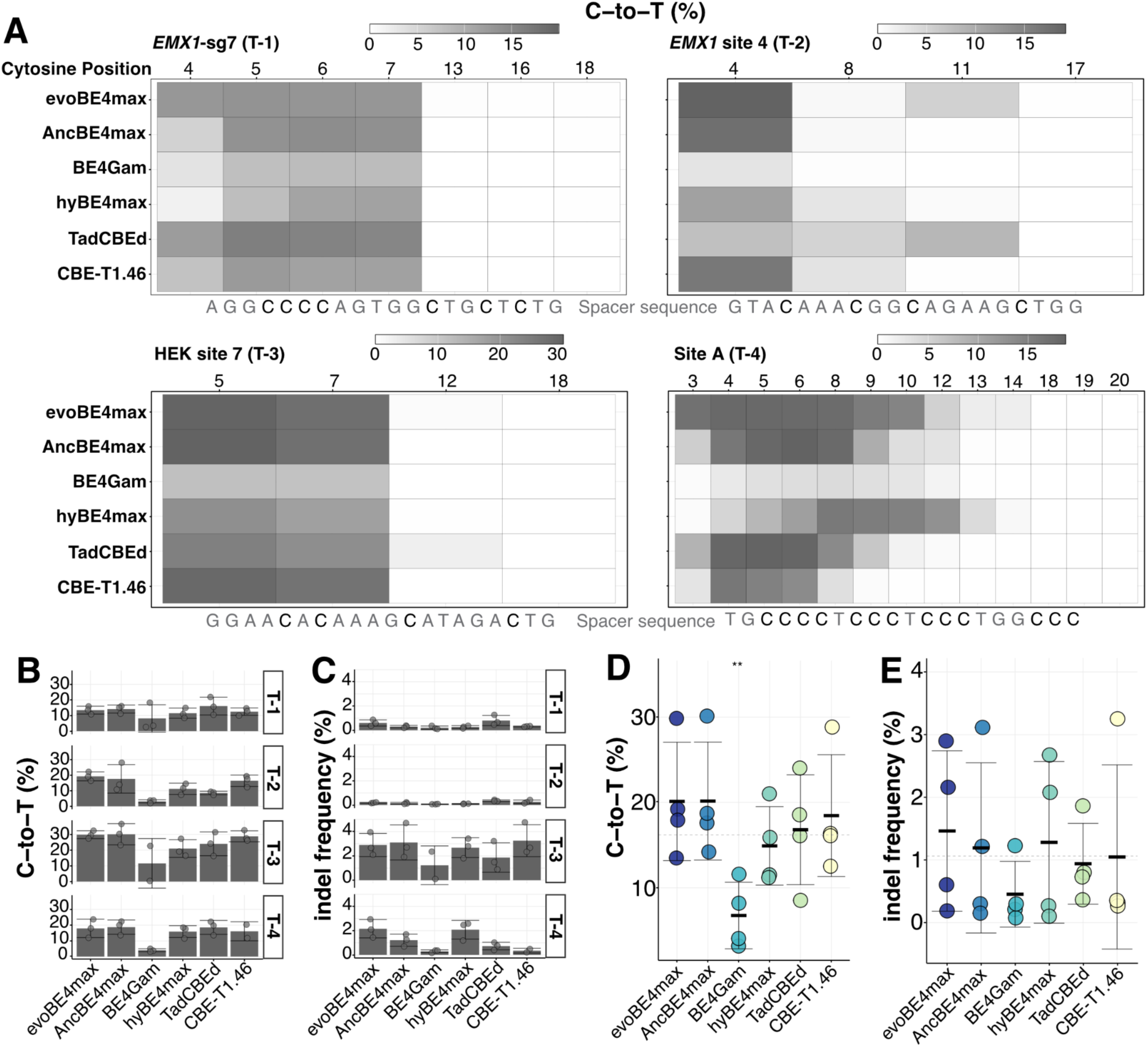
Benchmarking CBE variants in HEK293T cells. (A) C-to-T editing efficiency along the protospacer at four target sites, determined by amplicon deep sequencing. Heatmaps show the mean of three independent replicates. Protospacer sequences are depicted in grey below each heatmap. (**B-C**) Peak C-to-T editing (**B**) and indel frequency (**C**) of each CBE variant at the four target sites. Peak editing was defined as the highest C-to-T conversion rate along the protospacer for each sample. (**D-E**) Summary of peak C-to-T editing (**D**) and indel frequency (**E**) across all four loci. Each data point represents a single target site. The base mean (dotted line) represents the grand mean of peak editing across all six CBE variants. A one-sample *t*-test (two-tailed) was performed for each editor against the base mean; only BE4-Gam differed significantly (\*\**p*=.006) in (**D**). Data are represented as mean ± SD (n=3 independent replicates; n=2 for BE4-Gam at HEK site 7 and CBE-T1.46 at Site A). CBE, cytosine base editor.

Across the four target sites, evoBE4max demonstrated the broadest editing window (range of protospacer positions exhibiting at least half-maximal peak editing efficiency (Thuronyi *et al*., 2019)), with editing efficiencies of 13.0-29.8% across positions C3-10 at all four loci (Figure 1A). AncBE4max and CBE-T1.46 showed comparable efficiencies within narrower windows (C4-9 and C4-7, respectively); TadCBEd performed similarly, with modestly extended activity at peripheral positions; BE4-Gam showed substantially lower efficiencies across all sites; and hyBE4max did not reproduce its previously reported improvements over evoBE4max (Zhang *et al*., 2020). The level of indels remained below 1% for all editors at the *EMX1-*sg7 and *EMX1* site 4 loci but exceeded 2% for evoBE4max, AncBE4max, hyBE4max, and CBE-T1.46 at the HEK site 7 locus and was around 2% for evoBE4max and hyBE4max at the Site A locus (Figures 1A and 1B). EvoBE4max and AncBE4max achieved near-identical average peak efficiencies (20.1% and 20.2%) and indel rates (roughly 1.4%), but evoBE4max’s broader editing window and capacity to edit GC contexts made it our primary choice. Because evoBE4max shows reduced activity at AC dinucleotides (C8, *EMX1* site 4), in line with previous reports (Arbab et al., 2020; Cornean et al., 2022; Huang et al., 2021; Thuronyi *et al*., 2019), we designated CBE-T1.46, which lacks evident sequence context bias, as a backup editor (Figure 1C-E).

### Target gene sgRNA selection and validation

To identify sgRNAs capable of effectively disrupting *TPC1*, *TPC2*, and *OCaR2* gene function through PTCs or by interfering with SD or SA sites (Figure 2A), we selected the top three predicted sgRNA candidates for each of the three gene sequences for *in vitro* testing. The selection was based on the results of two sgRNA prediction tools (ACEofBASEs (Cornean *et al*., 2022), and IDT CRISPR-Cas9 guide RNA design checker), which ranked candidates according to editing efficiency, editing window, sequence context preferences, and the *in silico-*predicted number of off-target sites. To enrich for edited cells, we adapted the TREE reporter system (Standage-Beier *et al*., 2019), in which successful co-transfection of all editing components converts a BFP reporter to GFP, enabling FACS-based selection of edited populations (Figure 2B). To prioritise sgRNAs before committing to resource-intensive hiPSC editing, we first evaluated the peak editing efficiency of evoBE4max with nine sgRNAs targeting our candidate genes using the TREE reporter assay in HEK293T cells, followed by amplicon deep sequencing.

**Figure 2.**
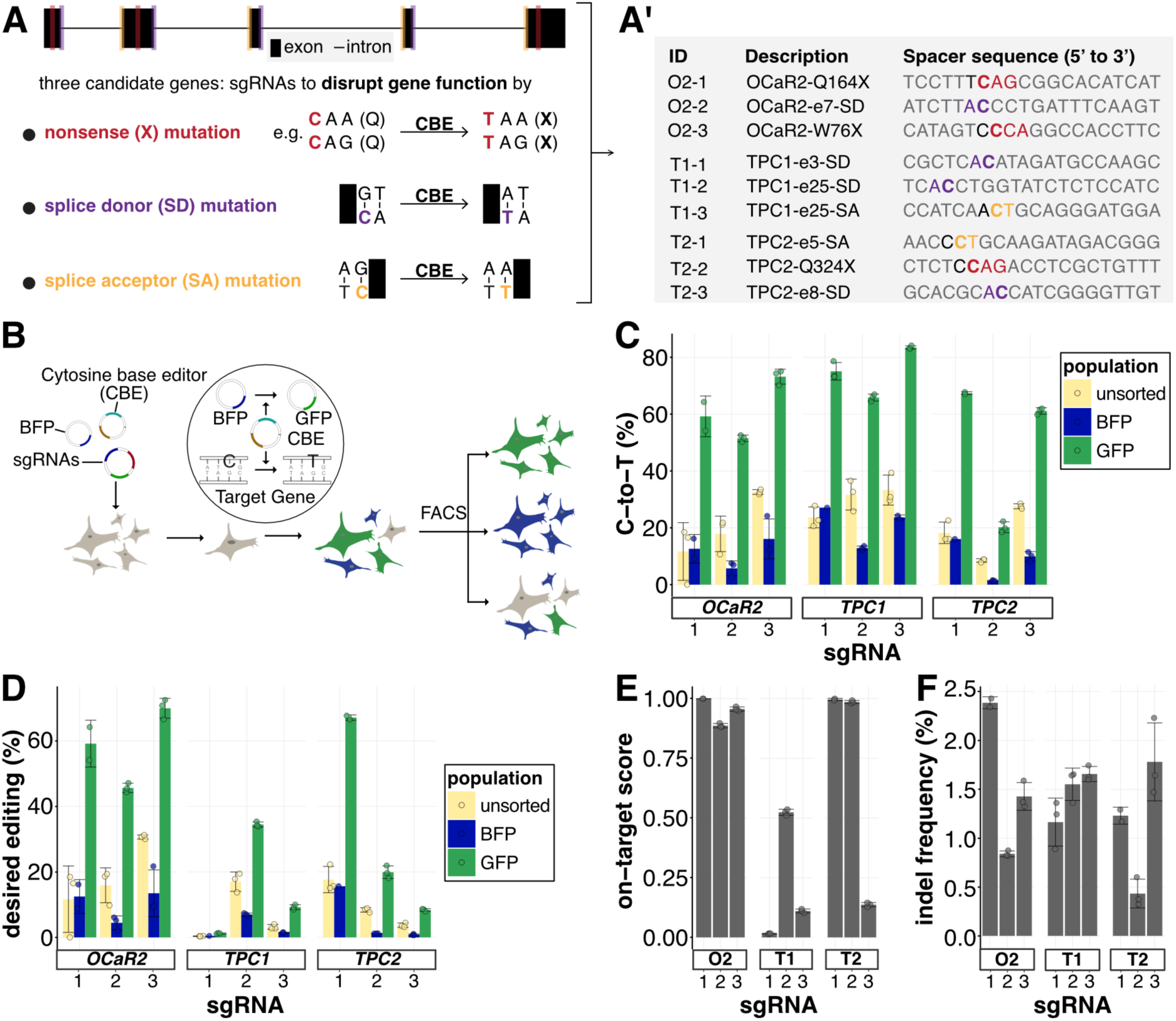
Validating candidate LoF sgRNAs in HEK293T cells. (A) Design of candidate sgRNAs for *OCaR2*, *TPC1*, and *TPC2* knockout via nonsense (X, dark red), splice donor (SD, purple), or splice acceptor (SA, orange) mutations. Along each protospacer, the target cytosine is shown in bold, the preceding nucleotide (sequence context) in black, and the remaining sequence in grey. (B) Transient reporter for editing enrichment (TREE) (Standage-Beier *et al*., 2019): a single C-to-T conversion in the co-transfected BFP reporter converts it to GFP, marking cells with active CBE. Three populations were analysed: unsorted (yellow), BFP-positive (blue), and GFP-positive (green). (C) Peak editing efficiency (highest C-to-T conversion within the protospacer). (D) Desired editing (editing at the target cytosine irrespective of bystander edits). (E) On-target score (ratio of desired to peak editing), shown for GFP-positive cells only. (F) Indel frequency, shown for GFP-positive cells only. Data are represented as mean ± SD (n=3 per population, except: n=2 for O2-1 BFP and GFP, T1-3 BFP, T2-1 GFP; n = 1 for T1-1 BFP, T2-1 BFP). BFP, blue fluorescent protein; O2, *OCaR2*; T1, *TPC1*; T2, *TPC2*. LoF, loss-of-function. See also Figure S1.

The top performing sgRNAs in the GFP-positive cell population were sgRNA O2-3 at the *OCaR2* locus, sgRNA T1-3 at the *TPC1* locus, and sgRNA T2-1 at the *TPC2* locus, achieving 73.2 ± 2.6% (at *C7),* 83.5 ± 0.6% (at *C5),* and 67.4 ± 0.5% (at *C4*) peak editing, respectively (Figure 2C). Across all sgRNAs, TREE enrichment increased peak editing efficiency 2.6- to 4.5-fold over unsorted and BFP-positive controls, respectively (Figure 2C). Because our LoF objective requires editing at a specific target cytosine, we also quantified desired editing, the fraction of reads carrying the disruptive C-to-T change regardless of bystander edits. Desired editing efficiencies closely tracked peak efficiencies for most sgRNAs (Figure 2D, Figure S1A), with notable exceptions discussed below. Interestingly, the on-target editing efficiency was significantly lower than the maximum editing efficiency for all three sgRNA candidates at the *TPC1* locus. The highest editing efficiency at the target nucleotide was observed with sgRNA T1-2, reaching 34.5 ± 0.8% in the GFP-positive population, possibly due to the disfavourable AC sequence context (Figure 2A’, Figure S1A). We also observed this decrease in desired editing efficiency for sgRNA T2-3 and, to a lesser extent, in sgRNA O2-2. Consequently, these sgRNAs showed a decreased “on-target score”, indicating the ratio between desired editing and the peak editing efficiency (Figure 2E). Throughout, the indel frequency remained below 2%, except for sgRNA O2-1 (Figure 2F, Figure S1C).

Given the high desired editing efficiency and low indel frequencies, we proceeded with sgRNAs O2-3, T1-2, and T2-1. We next validated the three selected sgRNAs in hiPSCs using the same TREE enrichment workflow. Peak editing efficiencies in the GFP-positive population were comparable to or higher than in HEK293T cells: 73.0 ± 0.1% for O2-3, 73.1 ± 16.0% for T1-2, and 82.6 ± 13.5% for T2-1 (Figure 3A). On-target editing rates followed the same pattern, with T1-2 remaining the lowest at 31.0 ± 14.5% (Figure 3B-C; Figure S1B). Notably, the enrichment effect was even more pronounced in hiPSCs, reaching up to 8.8-fold over unsorted cells, suggesting that TREE effectively compensates for the lower transfection and expression efficiencies typical of hiPSCs. Indel rates remained below 2% (Figure 3D, Figure S1D). Taken together, our two-stage pre-screening strategy, comprising systematic CBE benchmarking, sgRNA ranking in HEK293T cells, and TREE-based enrichment, yielded highly efficient editing at all three target loci in hiPSCs, establishing a validated toolkit for clonal derivation.

**Figure 3.**
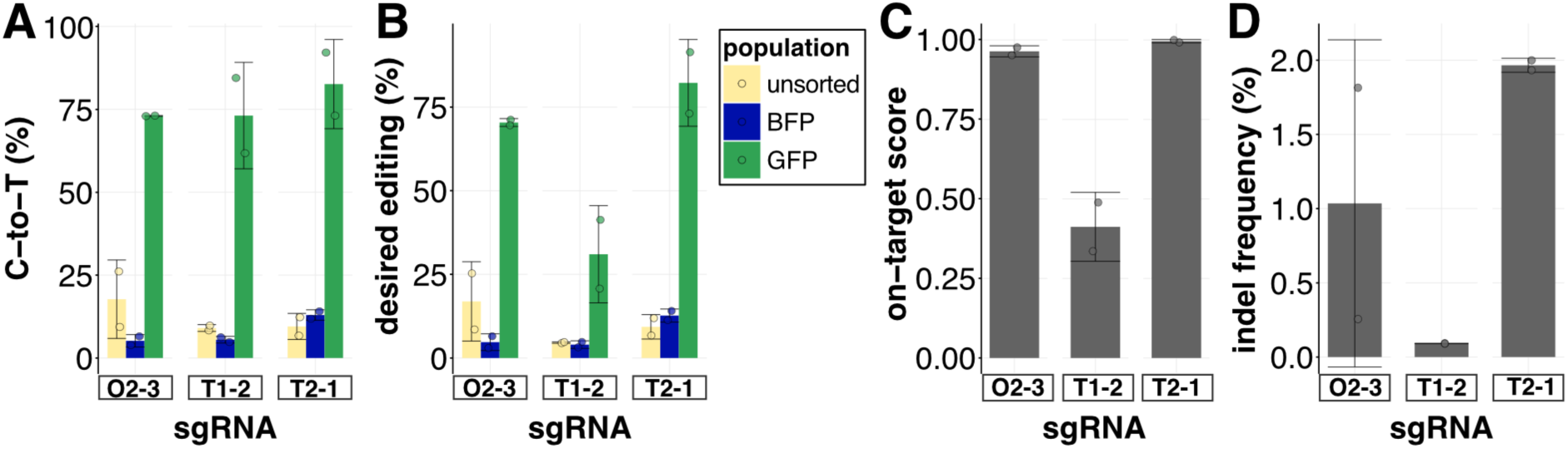
Validating LoF sgRNAs in hiPSCs. (**A-B**) One sgRNA per locus was selected and applied to hiPSCs. After FACS, (**A**) peak editing and (**B**) editing at the target cytosine were determined by amplicon deep sequencing for unsorted (yellow), BFP-positive (blue), and GFP-positive (green) cells. (**C-D**) On-target score (**C**) and indel frequency (**D**) for GFP-positive cells. Data are represented as mean ± SD (n=2 per population). LoF, loss-of-function. See also Figure S1.

### Generation of homozygous sKO and dKO hiPSC clones

Although multiplex CRISPR-Cas9 editing in hiPSCs has been achieved, efficient workflows for streamlined KO generation and comprehensive line validation remain limited (Brookhouser *et al*., 2020; Kim et al., 2025). Here, we have developed a simple and effective workflow for producing homozygous single-, double-, and triple-KO (sKO, dKO, and tKO) hiPSC lines by using an adapted version of the TREE assay (Standage-Beier *et al*., 2019; Tekel et al., 2021) and selecting suitable CBE variants and sgRNA candidates (Figure 4A). To maximise clonal survival, our protocol differs from previous workflows in three respects: (1) GFP-positive cells were first replated as a population before clonal isolation by limiting dilution, rather than direct single-cell sorting; (2) clonal expansion used a clump-promoting passaging solution instead of single-cell dissociation with Accutase; and (3) a specialised hiPSC survival supplement was added at three critical steps: before transfection, during the recovery phase following FACS, and in the final step of limiting dilution (Figure 4A). During the post-FACS recovery phase, we assessed bulk on-target editing using Sanger sequencing to triage populations prior to clonal isolation (Figure 4B).

**Figure 4.**
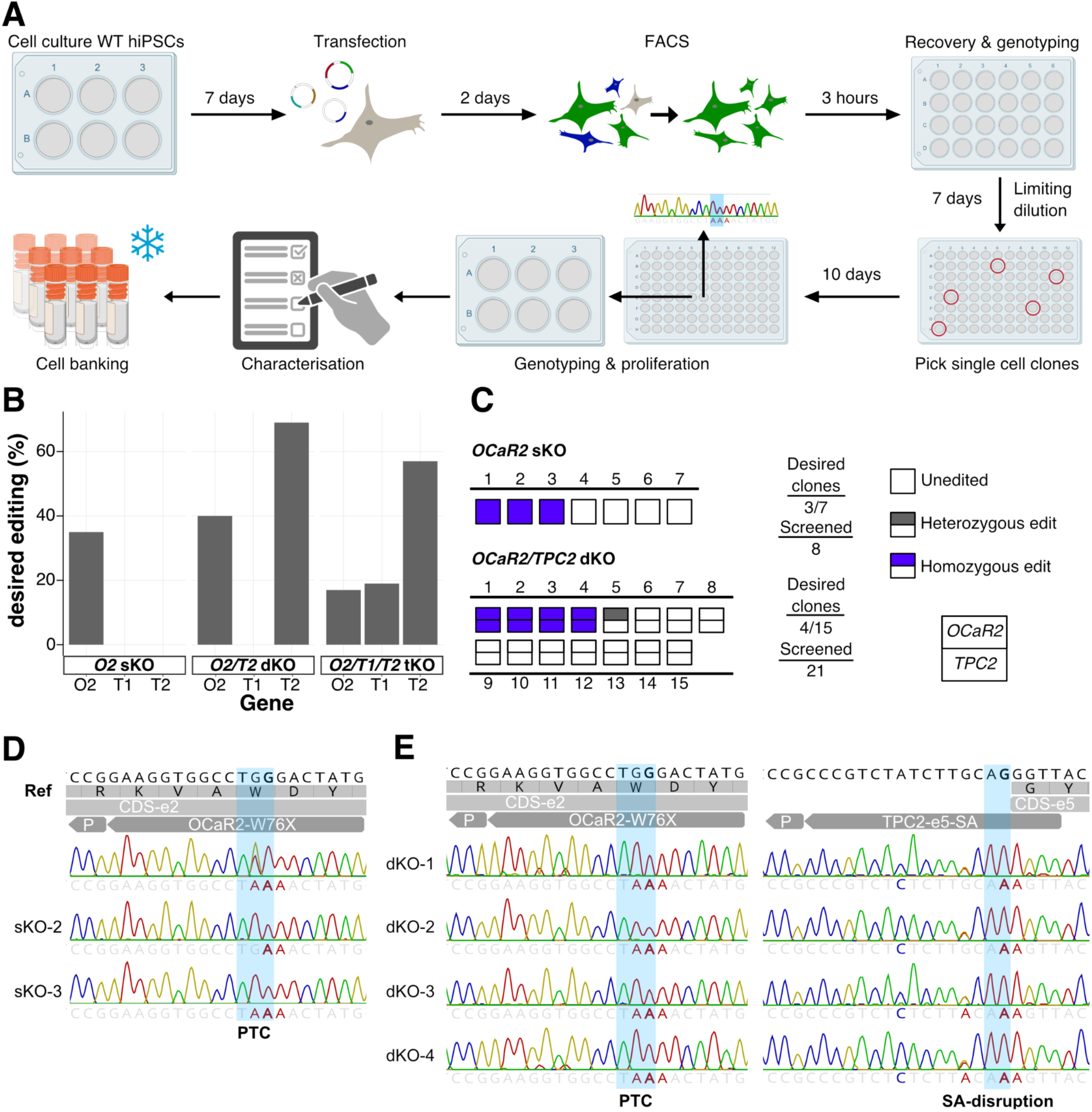
Generation of homozygous *OCaR2* sKO and *OCaR2*/*TPC2* dKO clones. (A) Workflow overview: after TREE transfection and FACS, GFP-positive cells are replated together for recovery. A sample of the bulk GFP-positive population is analysed by Sanger sequencing to assess editing efficiency. Limiting dilution is then performed to obtain single-cell clones, which are genotyped by Sanger sequencing during expansion. Selected KO clones underwent systematic characterisation and were cryopreserved. (B) Editing at the target cytosines in bulk GFP-positive cells after FACS (single measurement). (C) Overview of analysed *OCaR2* sKO and *OCaR2*/*TPC2* dKO clones: homozygous edits are shown in blue, heterozygous in grey, and wild-type in white. For dKO clones, each square is divided into two rectangles (upper: *OCaR2*; lower: *TPC2*). “Screened” indicates all populations analysed by Sanger sequencing; “desired clones” includes only clones confirmed to originate from a single cell. (**D-E**) Sanger sequencing chromatograms of identified homozygous *OCaR2* sKO (**D**) and *OCaR2/TPC2* dKO (**E**) clones. The wild-type protospacer sequence and corresponding amino acids are shown 5ʹ to 3ʹ at the top; with the coding sequence, exon number, sgRNA, and PAM beneath. Target nucleotides are highlighted in blue with the intended mutation indicated below. A single peak indicates homozygosity; two peaks indicate heterozygosity. Nucleotides differing from the reference are marked in colour. PTC, premature termination codon; SA, splice acceptor; CDS, coding sequence; e, exon; O2, *OCaR2*; T2, *TPC2*. Panel **A** was partially created with BioRender.com. See also Figures S2, S3, and S7 and Table S1.

Before proceeding to clonal isolation, we replated the FACS-sorted GFP-positive cells and expanded them for two to three passages to allow recovery (Figure 4A). Sanger sequencing of this recovered bulk population revealed that editing efficiencies had decreased relative to those measured directly after FACS in the initial validation experiments (Figure 3B): from 70% to 17-40% for *OCaR2*, from 31% to 19% for *TPC1*, and from 82% to 57-69% for *TPC2* across sKO, dKO, and tKO transfections (Figure 4B). This reduction suggests that a proportion of edited cells either underwent cell death or were outcompeted during recovery by unedited cells within the GFP-positive population. We continued with the GFP-positive populations, with editing efficiencies of 35% for single *OCaR2* edits and 40% and 69% for dual edits of *OCaR2* and *TPC2*, respectively (Figure 4B). We performed single-cell limiting dilution and split the surviving clones onto two 96-well plates - one for genotype validation and the other for clonal expansion. Clones harbouring the desired mutations were further characterised, expanded and cryopreserved (Figure 4A). We screened eight clones for *OCaR2* sKO and 21 clones for *OCaR2/TPC2* dKO using targeted Sanger sequencing, discarding populations that indicated non-clonal origins based on zygosity (Figure 4C). Three of the seven remaining *OCaR2* sKO candidates carried the homozygous LoF mutation p.W76X (Figure 4D). We kept one clone without editing at the *OCaR2* locus as a control line for subsequent QC assays (“deaminase control”, Deam ctr). For the *OCaR2*/*TPC2* dKO, four of the 15 remaining clones carried both the homozygous *OCaR2* p.W76X mutation and the homozygous *TPC2 SA-exon5* AG>AA mutation (Figure 4E). Notably, *OCaR2* sKO clones were exclusively either wild-type or homozygous mutants, with no heterozygous clones detected. Similarly, *OCaR2*/*TPC2* dKO mutations were mostly homozygous, with only one clone harbouring a heterozygous mutation (Figure 4C).

While the *OCaR2* sKO and *OCaR2*/*TPC2* dKO lines were generated in a single round of our workflow, retrieving *OCaR2*/*TPC1*/*TPC2* tKO clones proved more difficult. Our initial attempt showed low bulk editing efficiencies of 17%, 19%, and 57% for *OCaR2*, *TPC1*, and *TPC2*, respectively (Figure 4B). Based on observed editing efficiencies and clonal yields, including the predominantly bimodal (wild-type or homozygous) genotype distribution suggesting co-dependent allelic editing, we estimated that obtaining a tKO clone under these conditions would require screening an impractical number of clones (725-4503 wells; Exp. Procedures and Supplementary Table S3), prompting us to re-evaluate the editor and sgRNA for *TPC1*. Taken together, evoBE4max enabled efficient isolation of *OCaR2* sKO and *OCaR2*/*TPC2* dKO clones, but its reduced activity at the *TPC1* AC context necessitated optimisation for tKO generation.

### CBE-T1.46 enables triple-homozygous KO hiPSC line generation

To validate the overall efficacy of CBE-T1.46 in hiPSCs, we compared it with evoBE4max at HEK site 7, without TREE enrichment, where both editors achieved comparable peak efficiencies (42-44%) (Figure 5A). Based on our previous observation that GFP-positive cells following FACS sorting may exhibit a high level of subsequent cell death, we reduced the inputs of plasmid DNA and lipofection reagent (Exp. Procedures and Supplementary Table S2). Re-screening *TPC1* sgRNAs with CBE-T1.46 revealed that sgRNA T1-1 outperformed the previously selected T1-2 (53% vs 25% desired editing; Figure 5B), presumably due to the unbiased sequence preference of CBE-T1.46. Therefore, we repeated the transfection for the *OCaR2/TPC1/TPC2* tKO with sgRNA T1-1 and CBE-T1.46, resulting in editing efficiencies of 48%, 40%, and 71%, respectively (Figure 5C). These results are comparable to evoBE4max for *OCaR2* and *TPC2*. Subsequently, we screened 29 clones by targeted Sanger sequencing to identify the *OCaR2/TPC1/TPC2* tKO. We eliminated 10 clones due to zygosity ambiguities and identified one of the remaining 19 that possessed triple-homozygous LoF mutations: *OCaR2* p.W76X, *TPC1 SD-exon3* GT>AT*, and TPC2 SA-exon5* AG>AA (Figure 5D).

**Figure 5.**
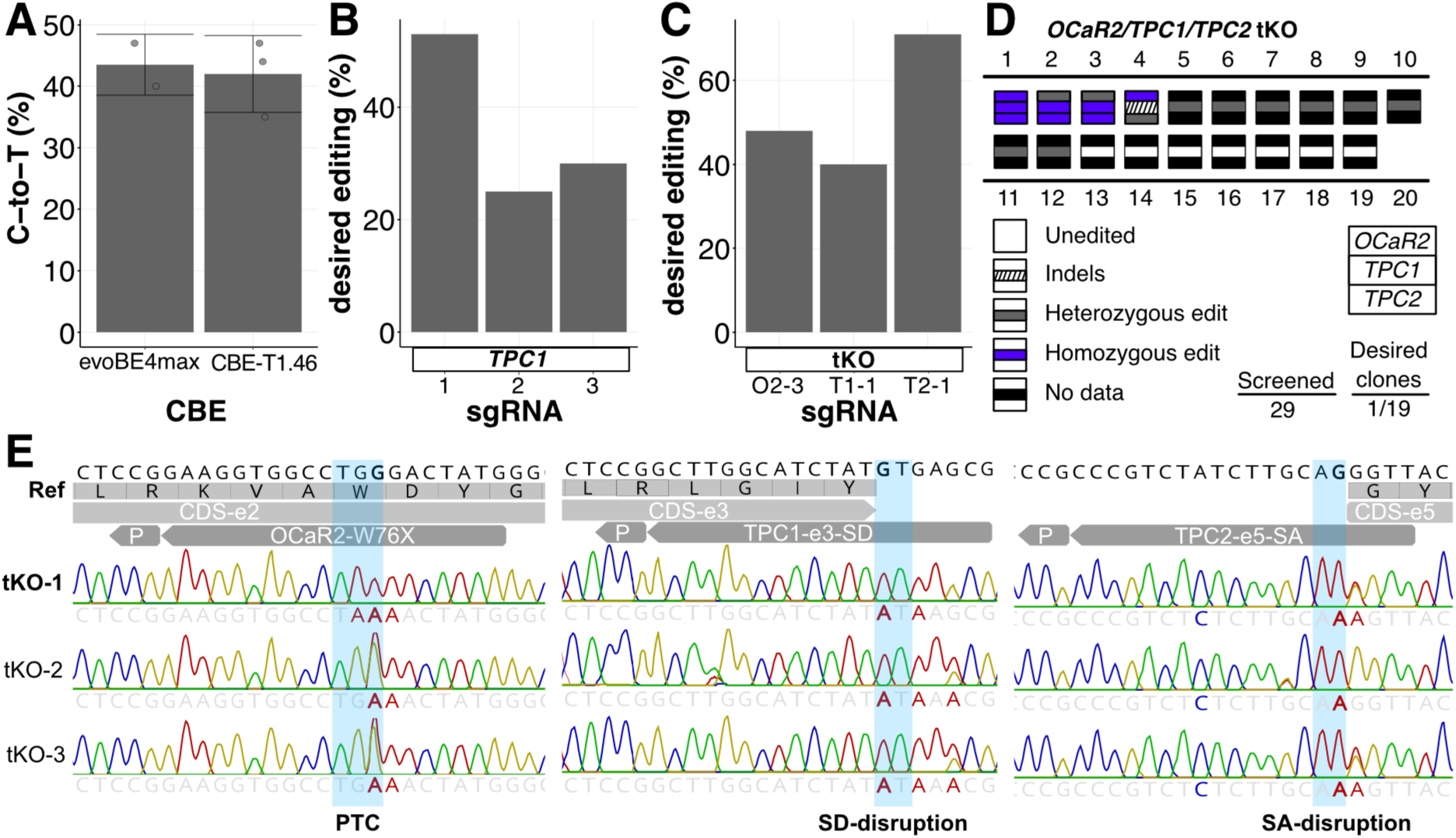
Troubleshooting *TPC1* editing in hiPSCs and generation of the *OCaR2*/*TPC1*/*TPC2* tKO. (A) Comparison of peak editing between evoBE4max (n=2) and CBE-T1.46 (n=3) at HEK site 7, determined by Sanger sequencing. Data are represented as mean ± SD. (B) *TPC1* sgRNAs retested in hiPSCs with CBE-T1.46; editing at the target cytosine is shown (single measurement, determined by Sanger sequencing). (C) Bulk on-target editing efficiency after simultaneous transfection of all three sgRNAs and FACS (single measurement, determined by Sanger sequencing). (D) Overview of analysed *OCaR2/TPC1/TPC2* tKO clones. Each square is divided into three rectangles (top to bottom: *OCaR2*, *TPC1*, *TPC2*): white, wild-type; grey, heterozygous; blue, homozygous; hatched, indel; black, no data. “Screened” indicates all populations analysed; “desired clones” includes only clones confirmed to originate from a single cell. (E) Sanger sequencing chromatograms of the identified tKO clone. Layout as in Figure 4D-E. PTC, premature termination codon; SA, splice acceptor; SD, splice donor; CDS, coding sequence; e, exon. See also Figures S2 and S7 and Table S1.

To reduce costs and save time, we initially analysed the *TPC1* locus and proceeded only with clones that showed a homozygous mutation at that locus. Compared to the previously created *OCaR2* sKO and *OCaR2*/*TPC2* dKO, the genotypes of the clones we analysed were more varied, with many heterozygous mutations, particularly in *TPC1*. In addition to the homozygous *OCaR2*/*TPC1*/*TPC2* tKO, we cryopreserved two clones that carried homozygous mutations in *TPC1* and *TPC2*, along with a heterozygous mutation at *OCaR2* (Figure 5E). By combining our enhanced transfection protocol with a gentle single-cell clone expansion using the backup editor CBE-T1.46, we successfully created a homozygous *OCaR2/TPC1/TPC2* tKO in just one attempt. Next, we assessed whether our selected hiPSC clones maintained pluripotency and lacked unwanted copy-number variations or off-target edits in their genome.

### CBE-mediated KO hiPSC lines retain pluripotency

To assess whether the KO clone generation workflow affected the undifferentiated hiPSC state, we monitored colony morphology and growth, and quantified established undifferentiated-state markers by RT-qPCR and immunofluorescence (Supplementary Figure S2). For downstream experiments, we considered clones suitable when they (1) retained typical hiPSC colony morphology and growth (Supplementary Figure S2A), and (2) showed concordant expression of the undifferentiated-state marker triad *OCT4*/*POU5F1*, *SOX2* and *NANOG* together with robust OCT4 and SSEA4 staining comparable to control cultures (Supplementary Figure S2B-C). While the majority of clones met these criteria, *OCaR2* sKO clone 3 and *OCaR2*/*TPC2* dKO clone 4 displayed aberrant, non-hiPSC-like morphology and a coordinated reduction of *OCT4*/*POU5F1*, *SOX2* and *NANOG* transcripts to <25% of wild-type levels, accompanied by reduced marker staining (Supplementary Figure S2B-C). Based on these features, these clones were excluded from further validation and phenotypic analyses. We next evaluated differentiation potential by directed trilineage differentiation in a representative subset of mutant clones (Supplementary Figure S2D-F). All tested clones generated derivatives expressing markers characteristic of endoderm, mesoderm, and ectoderm as assessed by RT-qPCR and immunofluorescence. The dKO clone showed reduced *TBXT* induction but retained functional mesodermal differentiation capacity as evidenced by cardiomyocyte differentiation (Supplementary Figures S2E and S7F). Importantly, the deaminase control, which was exposed to an active CBE but showed no on-target editing, displayed undifferentiated-state marker expression, a normal karyotype, and trilineage differentiation results similar to wild-type cells. This suggests that transfection, FACS-based enrichment, and clonal expansion did not significantly affect pluripotency at both the marker and functional levels (Supplementary Figure S2, Supplementary Table S1).

### Whole-genome sequencing reveals minimal guide-directed off-target editing

To assess genome integrity, we performed SNP-array karyotyping across edited clones and generated whole-genome sequencing (WGS) datasets for a subset of lines. SNP-array analysis did not reveal major copy number variants (CNVs), with the exception of a 475 kbp duplication on chromosome 2p in *OCaR2* sKO clone 1 and a 1051 kbp duplication on chromosome 9 in *OCaR2/TPC2* dKO clone 3 (Supplementary Figure S3, Supplementary Table S1). Next, we assessed genome-wide off-target editing by evoBE4max. We performed WGS of the *OCaR2* sKO (O2-sKO) clone 1 and *OCaR2*/*TPC2* dKO (O2T2-dKO) clone 1 alongside the parental wild-type hiPSC line. These two clones were selected because both were edited with evoBE4max, whose rAPOBEC1-derived deaminase induces measurable genome-wide C-to-T mutagenesis (Doman et al., 2020; Yu et al., 2020; Zuo et al., 2019), whereas the *OCaR2*/*TPC1*/*TPC2* tKO clone was generated using CBE-T1.46, for which WGS has already demonstrated minimal stochastic deamination (Lam *et al*., 2023). Somatic variants were called against the parental line and filtered stringently (Kim *et al*., 2025), then classified into four tiers based on overlap with Cas-OFFinder-predicted off-target sites (Bae et al., 2014), and consistency with CBE activity (Exp. Procedures; Figure 6, Supplementary Figures S4-6).

**Figure 6.**
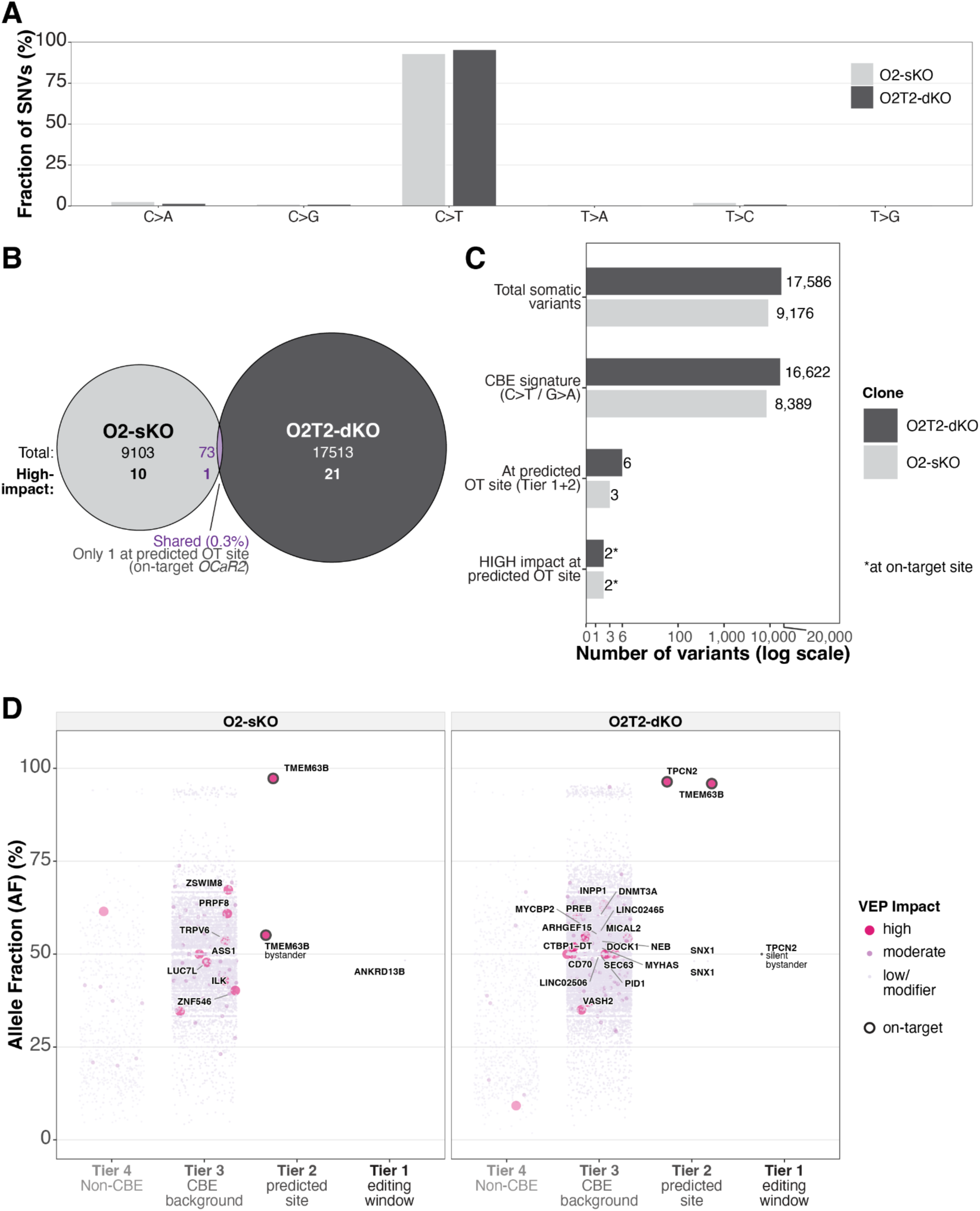
Whole-genome sequencing reveals minimal guide-directed off-target editing in evoBE4max-edited hiPSC clones. (A) Mutation spectrum showing the six substitution classes for the O2-sKO and O2T2-dKO clones, showing C>T/G>A dominance at 92.9% and 95.4%, respectively. (B) Venn diagram of variant overlap between clones. Only 73 of more than 26,000 combined variants (0.3%) were shared; the single shared variant at a predicted off-target site mapped to the on-target *OCaR2* locus. Numbers in parentheses indicate HIGH-impact variants unique to each clone. (C) Variant filtering funnel (pseudo-log scale) showing progressive reduction from total somatic variants to HIGH-impact variants at predicted off-target sites. Only 3 (O2-sKO) and 6 (O2T2-dKO) variants mapped to any of 420 predicted off-target windows; no HIGH-impact coding variant was detected at a genuine off-target site. (D) Variant landscape plots displaying allele fraction versus tier classification for each clone. Variants are coloured by VEP functional impact (HIGH, red; MODERATE, purple; LOW/MODIFIER, light purple). Gene labels indicate Tier 1+2 variants and clonal (AF≥0.4) HIGH-impact Tier 3 variants. On-target edits at *TMEM63B* (*OCaR2*) and *TPCN2* (*TPC2*) are marked. The sole genuine off-target Tier 1 variant (chr17:29,607,330, O2-sKO) is non-coding; the O2T2-dKO Tier 1 variant maps to the on-target TPC2 locus. n=1 clone per genotype. See also Figures S4-S6.

We detected 9,176 and 17,586 *de novo* somatic variants in the O2-sKO and O2T2-dKO clones, respectively. The single-nucleotide variant (SNV) spectrum was overwhelmingly dominated by C>T/G>A transitions (92.9% and 95.4%) (Figure 6A), with C-to-T odds ratios of 65.7 and 104.5 relative to a uniform substitution model (Supplementary Figure S5B). Trinucleotide context analysis revealed enrichment at TC (approximately 40%) and CC (approximately 41-42%) dinucleotide motifs, with depletion at both GC (<3%) and AC contexts (1.1% and 0.8%) (Supplementary Figure S4C), the latter mirroring the disfavoured AC dinucleotide context we observed for evoBE4max at on-target sites during editor benchmarking (Figure 1A). Approximately 84% of background C>T variants were clonal (AF≥0.4), indicating deamination at or near the time of editing (Supplementary Figure S5A). Only 73 variants (0.3%) were shared between the two independently edited clones, of which only a single variant mapped to a predicted off-target site - at the on-target *OCaR2* locus itself (Figure 6B). Together, these features confirm that the *de novo* variants reflect stochastic, guide-independent cytosine deamination by rAPOBEC1 rather than culture-associated mutagenesis, at levels comparable to those reported earlier (Doman *et al*., 2020; Gaudelli et al., 2020; Kim et al., 2017; McGrath et al., 2019; Young et al., 2012; Zuo *et al*., 2019). The O2T2-dKO clone carried approximately twice the *de novo* burden of the O2-sKO clone despite identical transfection conditions, FACS timing, and passage history. The number of variants at predicted off-target sites scaled with the number of guides (3 versus 6 for one versus two guides; Figure 6C), consistent with additive guide-directed risk. By contrast, the two-fold increase in guide-independent Tier 3 background (8,388 versus 16,621) is better explained by cell-to-cell variability in CBE expression: cells achieving simultaneous editing at two loci are likely enriched for higher deaminase activity, resulting in a proportionally larger stochastic C>T background (Figure 6A).

Progressive filtering dramatically reduced the variant burden (Figure 6C). Of 420 predicted off-target windows, only 3 and 6 variants in the O2-sKO and O2T2-dKO clones mapped to any predicted site, most at the intended on-target loci. Each clone harboured one Tier 1 variant: in the O2-sKO clone, a heterozygous non-coding G>A at chr17:29,607,330 (3-mismatch site for guide O2-3), the sole genuine guide-directed off-target SNV detected across both clones (Supplementary Figure S6A); in the O2T2-dKO clone, a G>A within the on-target *TPC2* locus itself, representing intended editing. Three additional off-target variants at 3-mismatch sites were identified in the O2T2-dKO clone, all non-coding (Supplementary Figure S6B-C). Critically, no Tier 1 or Tier 2 variant in either clone resulted in a coding change at a genuine off-target site (Figure 6B-D). An orthogonal guide homology analysis confirmed that all 66 HIGH-impact variants outside on-target loci had 8 or more mismatches to either guide, far beyond the range of Cas9-mediated editing (Supplementary Figure S4B). VEP annotation identified 29 and 37 HIGH-impact variants (stop-gained, frameshift, splice-site) in the O2-sKO and O2T2-dKO clones, all Tier 3 or Tier 4 background events (Figure 6D). Two ClinVar-flagged variants in the O2-sKO clone (an *ASS1* splice-acceptor variant and a low-frequency *ADAMTS13* missense variant) were both Tier 3 with 9 or more mismatches to either guide, confirming stochastic origin (Supplementary Figure S5C). No ClinVar pathogenic variants were detected in the O2T2-dKO clone. Manta and CNVKit analysis detected no structural variants or copy number changes at any predicted off-target site (Supplementary Figure S4A). Taken together, these analyses demonstrate that while evoBE4max introduces a characteristic stochastic CBE mutational background, guide-directed off-target editing is minimal, supporting the suitability of these clones for downstream functional studies.

### Molecular validation of LoF alleles across edited hiPSC lines

We designed validation assays matched to each edit type: transcript-level analysis for splice-site disruptions (*TPC1*, *TPC2*) and additional protein-level assessment for the PTC in *OCaR2*, where nonsense-mediated decay (NMD) can be incomplete (Khajavi et al., 2006; Sato and Singer, 2021) (Supplementary Figure S7A).

### *TPC2* (SA exon 5, AG→AA): reduced abundance of correctly spliced exon 5-containing transcript

To probe the impact of the exon 5 splice acceptor edit on *TPC2* RNA, we performed RT-qPCR using a primer pair amplifying across exon 5 to exon 6 (forward primer in exon 5; reverse primer in exon 6) (Supplementary Figure S7A-B). Across presumptive *TPC2* KO genotypes, exon 5 inclusion was consistently lower compared to wild-type, indicating a decreased abundance of the normally spliced transcript that includes exon 5. The splice acceptor mutation is expected to disrupt exon definition, is predicted to cause exon 5 skipping and to produce an in-frame deletion of 39 amino acids within the PIP2-binding domain (She et al., 2019). This outcome is consistent with a predicted LoF or strongly hypomorphic *TPC2* allele.

### *TPC1* (SD exon 3, GT→AT): increased intron retention and cDNA-level confirmation of aberrant splicing and a downstream PTC

Using a targeted intron-retention assay, we observed a marked increase in exon 3-intron 3 retention (roughly 8.0-fold relative to wild-type; Supplementary Figure S7C-D). Sanger sequencing of the corresponding cDNA amplicon confirmed disruption of the exon 3 splice donor and the presence of the predicted aberrant transcript carrying a downstream PTC (Supplementary Figure S7C, E). These results support that *TPC1* transcripts are predominantly mis-spliced, consistent with a predicted LoF allele.

### *OCaR2* (p.W76X): heterogeneous transcript reduction and marked protein depletion in differentiated cardiomyocytes

For *OCaR2* p.W76X, RT-qPCR showed reduced but heterogeneous transcript levels across edited lines, with many clones retaining ≥50% of wild-type *OCaR2* mRNA (Supplementary Figure S7A-B), consistent with partial NMD and/or NMD escape reported for PTC (Khajavi *et al*., 2006; Sato and Singer, 2021). We therefore assessed OCaR2 at the protein level in a context of robust expression by differentiating wild-type and *OCaR2/TPC2* dKO hiPSCs into atrial cardiomyocytes (aCMs). Quantitative immunofluorescence analysis (normalised to cardiac troponin T, cTnT) showed a ∼4-fold reduction of OCaR2 signal in dKO aCMs compared with wild-type (Supplementary Figure S7F-G), indicating a tendency for a depletion (*p*=.052) of OCaR2 protein in the edited line and supporting a predicted LoF *OCaR2* allele.

Across all genotypes, disruptive DNA edits produced concordant RNA and/or protein consequences, supporting predicted LoF alleles in *OCaR2*, *TPC1*, and *TPC2*.

### Extension of multiplex editing to *ORAI* Ca²⁺ channel paralogs

To test whether our CBE-based workflow extends beyond *OCaR2*/*TPC* targets, we applied evoBE4max to the *ORAI1*, *ORAI2* and *ORAI3* loci (Rubaiy, 2023) (Figure 7, Supplementary Figure S8). To streamline sgRNA selection, we pre-screened candidates in HEK293T cells using FACS enrichment of transfected cells, followed by EditR-based sequence deconvolution of Sanger traces (Kluesner et al., 2018) (Figure 7A; Figure S8A-D), thereby reducing the number of guides carried forward into hiPSC editing. Editing in HEK293T cells was consistently high across all nine sgRNAs, with desired editing at the target cytosine ranging from 46.7% to 98.7% (Figure 7A). Based on this pre-screen, we selected sgRNAs ORAI1-3, ORAI2-2, and ORAI3-1 for hiPSC experiments (Figure 7A), noting that ORAI2-2 maintained robust editing despite an AC dinucleotide context at the target site. Based on this pre-screen, we selected sgRNAs ORAI1-3, ORAI2-2, and ORAI3-1 for hiPSC experiments (Figure 7A), noting that ORAI2-2 maintained robust editing despite an AC dinucleotide context at the target site. We selected ORAI1-3 for its editing efficiency, comparable to that of ORAI1-2, but with a better bystander editing profile. In hiPSCs, transfection with single sgRNAs yielded high desired editing across loci (mean ∼82% per target; Figure 7B). Simultaneous delivery of all three sgRNAs also produced high editing efficiencies at each locus (88% for *ORAI1*, 83% for *ORAI2* and 90% for *ORAI3*; Figure 7C, Figure S8E-G). These results demonstrate that the sgRNA selection and bulk editing stages of our workflow transfer efficiently to additional gene families. Clonal isolation, genotype confirmation, and LoF validation for the *ORAI* targets remain to be completed.

**Figure 7.**
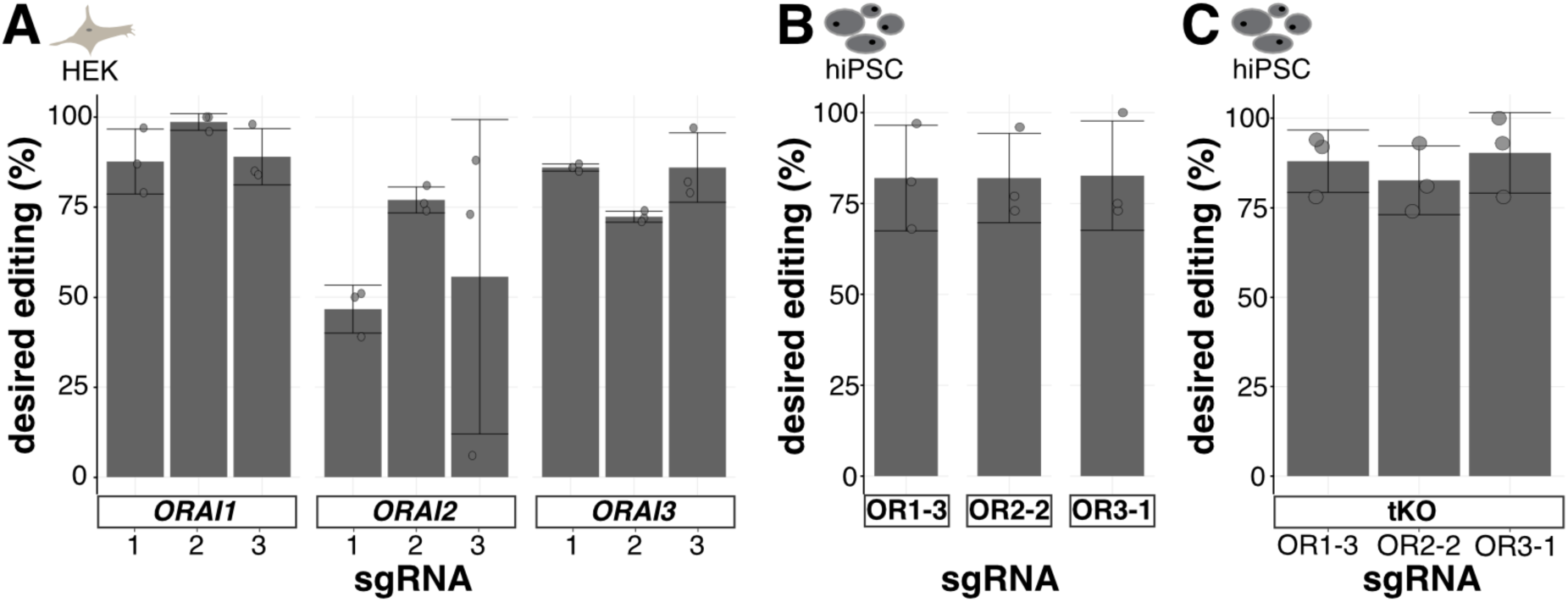
Application of the workflow to *ORAI* Ca²⁺ channel paralogs. (**A-B**) Editing at the target nucleotide in bulk GFP-positive cells after FACS for all *ORAI1-3* candidate sgRNAs in HEK293T cells (**A**) and for selected sgRNAs in hiPSCs (**B**). (**C**) On-target editing of bulk GFP-positive cells for the triple-KO after simultaneous transfection of all three sgRNAs and FACS. All data were obtained by Sanger sequencing. Data are represented as mean ± SD (n=3). See also Figure S8.

## DISCUSSION

We present a comprehensive, DSB-independent workflow for multiplex (≥2–3 loci) LoF editing in hiPSCs, combining systematic editor/sgRNA selection, transient reporter-based enrichment, and structured QC of pluripotency and genome integrity. Using cytosine base editing, we generated homozygous single-, double-, and triple-mutant hiPSC lines by inserting PTCs and disrupting splice sites, validated at the genotype level and through orthogonal transcript and protein assays. We also provide practical decision points for guide selection and realistic expectations for clone yield.

CBE-mediated PTCs and splice-site disruptions produce predictable LoF alleles that are more straightforward to confirm than frameshift-dependent indels (Kluesner *et al*., 2021). While previous pipelines have demonstrated scalable single-gene LoF engineering in hiPSCs using PTC-focused base editing with reporter enrichment and systematic QC (Zhang *et al*., 2024b), as well as multiplex base editing (Brookhouser *et al*., 2020), these efforts did not combine concurrent multi-locus editing with structured assessment of genome integrity. Our workflow extends these foundations to multiplex editing (single- through triple-KO) using both PTC and splice-site strategies, integrated with pluripotency and genome integrity QC and explicit guidance on clone yield - elements often under-reported yet critical for scalable application. Splice site editing provides an alternative when PTC introduction is infeasible or risks NMD escape (Khajavi *et al*., 2006; Sato and Singer, 2021) and enables a direct transcript-level readout that facilitates rapid clone triage in multiplex workflows.

An important practical insight is that the choice of editor and guide architecture is a primary factor influencing editing outcomes in hiPSCs. We observed robust performance with state-of-the-art CBEs across various target sites beyond the initial set of target genes, demonstrating the versatility of the editor-plus-enrichment approach. However, we also encountered locus-specific limitations that in one case necessitated switching editors. We therefore recommend a lightweight pre-screening approach, such as testing in HEK293T or bulk hiPSC populations, to rank sgRNAs and editors prior to clonal work. For guide selection, our data support focusing on (1) early coding exons when applying PTC strategies (Billon *et al*., 2017; Cuella-Martin et al., 2021; Hanna et al., 2021; Xu et al., 2021), (2) protospacers with fewer editable cytosines when multiple guides have similar efficiency to minimise bystander effects, and (3) target cytosines within the empirically effective activity window (mostly mid-protospacer) to maximise desired editing. Looking ahead, ongoing editor development is expected to enhance product purity and minimise unintended edits. This includes newer TadA-derived CBEs as well as emerging architectures that improve purity and reduce indels (Hu et al., 2026; Wu et al., 2026; Zhang et al., 2024a). Additionally, advanced splice-disruption strategies (e.g., synchronised SA+SD targeting) may increase the reliability of splice-based LoF designs, especially where single-site disruption produces heterogeneous results (Miskalis et al., 2024).

A key advantage of base editing is its independence from DSBs. CRISPR-Cas9 nuclease editing can induce p53-dependent toxicity in hiPSCs (Ihry *et al*., 2018), large deletions, chromosomal rearrangements, and copy-number changes at or near the cut site (Cullot et al., 2019; Hwang et al., 2025; Leibowitz et al., 2021; Liu et al., 2021), reducing viability and complicating downstream analyses; in iPSCs specifically, 33% of CRISPR/Cas9-edited clones harboured large on-target defects that escaped standard PCR-based screening (Simkin et al., 2022). However, “DSB-independent” does not mean there is no genotoxicity: base editing and prime editing can still produce on-target byproducts, such as indels and sometimes larger rearrangements (George et al., 2024; Hwang *et al*., 2025), with cellular stress responses observed in primary stem cell types (Fiumara et al., 2024). The appropriate approach is to focus on risk reduction rather than complete elimination, which is why genome integrity QC is an essential part of any clonal hiPSC engineering process.

In line with this, pluripotency and genome integrity were explicitly included as core criteria rather than afterthoughts, consistent with established best-practice recommendations for iPSC qualification (Hägg et al., 2026; Steeg et al., 2021). Our WGS analysis of evoBE4max-edited clones illustrates this risk-reduction principle in practice. While we detected thousands of *de novo* C>T/G>A variants consistent with stochastic rAPOBEC1-mediated deamination (Doman *et al*., 2020; Gaudelli *et al*., 2020; McGrath *et al*., 2019; Zuo *et al*., 2019), tiered classification revealed that guide-directed off-target editing was minimal: only one non-coding off-target SNV was identified at a predicted site across both sequenced clones, and no coding off-targets were detected. Of note, the total *de novo* burden approximately doubled in the dKO clone relative to the sKO clone despite identical transfection conditions, suggesting that cells achieving multiplex editing may be enriched for higher CBE expression and consequently greater stochastic deamination - a consideration relevant to any enrichment-based multiplex CBE workflow. The use of TadA-derived CBEs such as CBE-T1.46, which exhibit substantially lower guide-independent deamination (Lam *et al*., 2023), may mitigate this burden in future applications.

Several limitations should be noted. First, although we incorporated genome integrity as a key QC gate using SNP-array karyotyping across edited clones, WGS has so far been generated and analysed only for a subset of lines (control, sKO, and dKO), and additional WGS-based assessment across further clones will strengthen conclusions about genome-wide variation associated with multiplex base editing. Platform-specific differences in CNV detection sensitivity (e.g., a chr2p duplication detected by SNP-array but not WGS-based CNVKit) argue for using complementary approaches when assessing genome integrity. Second, our workflow was established in a single hiPSC genetic background; while we demonstrate portability across two independent triple-target configurations and multiple loci, extending the approach to additional donor backgrounds will be important for defining general applicability and identifying context-specific constraints. Third, we primarily validate gene disruption at the molecular level (by identifying disruptive genotypes with concordant transcript and/or protein consequences). While this supports the identification of predicted LoF alleles suitable for many applications, functional assays tailored to each target would be required to confirm complete loss of protein activity, particularly for alleles expected to be hypomorphic. Finally, clonal derivation remains a bottleneck for multiplex engineering in hiPSCs: enrichment and single-cell cloning impose stress and may select for fitter subclones, and limiting dilution can carry a residual risk of multi-cell colony origin; semi-automated or nanowell-based cloning workflows (Frank et al., 2023; Takahashi et al., 2025) and reporter-free enrichment strategies such as selective nuclease-induced purity enhancement (Fast et al., 2025), may further streamline multiplex workflows.

Taken together, this framework supports more predictable multiplex LoF engineering in hiPSCs while underscoring the importance of standardised QC as editing complexity increases.

## EXPERIMENTAL PROCEDURES

### Design and cloning of sgRNAs

sgRNAs were designed manually to enable the installation of LoF mutations (PTC, SD or SA mutations) and assessed for predicted editing outcomes using ACEofBASEs (Cornean *et al*., 2022), and the CRISPR-Cas9 guide RNA design checkers (IDT) with the “evoBE4max” BaseEditor pre-settings. For TREE-transfections, a BbsI-digested pDT-sgRNA backbone (pDT-sgRNA was a gift from Xiao Wang, Addgene plasmid #138271) and oligonucleotides (Sigma Aldrich, Merck) containing the sgRNA sequence with the following overhangs (5’-CACC(G)(N_1-20_)-3’ (forward) and 5’-AAAC(N_1-20_)(C)-3’ (reverse)) were cloned into plasmids as previously described (Standage-Beier *et al*., 2019). The sequences of the sgRNA plasmids were verified by Sanger sequencing and are listed in Supplementary Table S5.

### Cloning of CBE plasmids

For transfection in hiPSCs, a pEF1α backbone (from pEF1α-hMLH1dn, a gift from David Liu, Addgene Plasmid #174824) (Chen et al., 2021), and either evoBE4max (pBT281, a gift from David Liu, Addgene Plasmid #122611) or CBE-T1.46 (a gift from Nicole Gaudelli, Addgene Plasmid #193284) were PCR-amplified (Lam *et al*., 2023; Thuronyi *et al*., 2019), followed by Gibson assembly (key resources table, Supplementary Table S5). The whole sequences of both constructs were verified with the PlasmidEZ (Azenta/GeneWiz) full-plasmid sequencing service. EndoFree Plasmid Maxi Kit (Qiagen) was used for the preparation of the plasmids.

### Cell lines and cell culture

HEK293T cells and hiPSCs were cultivated as previously described (Ong et al., 2026). In brief, HEK293T cells were purchased from ATCC, and maintained in Dulbecco’s modified Eagle’s medium (DMEM; Gibco) supplemented with 10% FBS (Gibco). The hiPSC line SCVI15 (male), obtained from Greenstone Bioscience, Inc, was cultured on hESC-Qualified Matrix Matrigel-coated (Corning) 6-well plates (Sarstedt) in StemMACS PSC-Brew XF medium (Miltenyi Biotec). Beyond the information listed in the key resources table, the source provided no additional donor background or genomic characterisation information. Wild-type hiPSCs (Passage 29) were used for all transfections. During KO-hiPSC line generation and characterisation, cells were expanded for up to 24 passages or a total of 53 passages before cell banking.

### Transfection protocol

For CBE selection, transfections in HEK293T cells were performed as previously described (Ong *et al*., 2026). For TREE experiments, HEK293T cells were seeded on 12-well plates (130,000 cells/well) once they reached 70-80% confluency. After 24 hours, cells were transfected with 200 ng BFP and sgRNA plasmid, 600 ng CBE plasmid and 4 µl Lipofectamine 2000 (Invitrogen) per well. The transfection mix was added dropwise to the well. The medium was changed the next day. On the second day after transfection, cells were prepared for FACS. For hiPSCs, on the third day after the last passaging or when the hiPSCs reached more than 90% confluency, cells were seeded onto a 6-well plate for transfection (300,000 cells per well) using StemMACS Passaging Solution XF (Miltenyi Biotec). During seeding, 2.5 µM StemMACS Y-27632 (Miltenyi Biotec) was added until proceeding with the transfection. The transfection was started 24-36 hours after seeding, when the cells reached 50% confluency. Prior to transfection, the medium was changed to PSC-Brew XF medium supplemented with StemMACS PSC-Support XF (Miltenyi Biotec), an optimised supplement that improves human pluripotent stem cell performance under challenging culture conditions. Plasmid amounts per well of a 6-well plate were scaled according to the number of target loci (Supplementary Table S2), and Lipofectamine Stem (Invitrogen) was added. In all conditions, Opti-MEM (Gibco) was added to a total reaction volume of 330 µl. After 15 min of incubation, the reaction mix was applied dropwise to the cells. The medium was changed the following day, and on the second day after transfection, the cells were prepared for FACS.

### FACS procedure

HEK293T cells and hiPSCs were prepared for FACS as previously described (Ong *et al*., 2026). Sorting gates were set based on the respective single-colour control samples (BFP- or GFP-positive). FACS was performed with a BD FACSAria III at the Flow Cytometry & FACS Core Facility (FFCF), Zentrum für Molekulare Biologie der Universität Heidelberg (ZMBH). After FACS, the GFP-positive hiPSCs were collected in 500 µl StemMACS PSC-Brew XF medium supplemented with 2.5 µM Y-27632, 1% (v/v) Penicillin/Streptomycin (10,000 U/ml Pen, and 10,000 µg/ml Strep, Gibco) and 0.1% (v/v) Normocin (50 mg/ml, InvivoGen). The cell suspension was centrifuged for 5 min at 150 × *g*, and the supernatant was discarded. The cell pellet was resuspended in StemMACS PSC-Brew XF medium supplemented with 2.5 µM Y-27632, 1% Penicillin/ Streptomycin, 0.1% Normocin and StemMACS PSC-Support XF and replated in a well format appropriate to the number of cells (e.g. 30,000 cells in a 48-well plate). The cells were maintained in antibiotic-containing medium (1% Penicillin/Streptomycin, 0.1% Normocin) in StemMACS PSC-Support XF until the next passaging. During the next splitting, the cells were divided between two wells: one for assessment of bulk editing efficiencies and one for further expansion. For genome DNA extraction, the cells were incubated with a gDNA extraction buffer (10 mM Tris-HCl, pH 8.0; 0.05% SDS; 800 U/mL proteinase K (NEB)) at 37°C for 2 hours. Proteinase K was inactivated at 80°C for 30 min.

### Genotyping

The gDNA was diluted to 50 ng/µl with nuclease-free water. The PCR was performed with Q5 Hot Start Polymerase (NEB) according to the manufacturer’s instructions using 3 µl gDNA and following cycling program: 2 min at 98°C, 30-32 cycles of 30 sec at 98°C, annealing for 20 sec and 20 sec at 72°C, followed by 5 min at 72°C. Primers are listed in Supplementary Table S5. After agarose gel electrophoresis, amplicons were purified using the Monarch Gel extraction kit (NEB). Sanger sequencing was carried out by Eurofins Genomics as a service. Sanger sequence analysis was performed in Geneious Prime (2024.0.2), and the transition rates of base editing experiments were estimated using EditR (version 1.0.10) (Kluesner *et al*., 2018).

### Generation of hiPS single-cell clones

Following the replating after FACS, the cells were expanded stepwise to 12-well plates. When they reached 90% confluency, cells were counted and diluted to 25,000 cells/ml, then serially diluted (1:50, 1:17, 1:5) to a final concentration of 6 cells/ml (0.6 cells/well in 100 µl) before seeding into 96-well plates. The single-cell clones were seeded into 96-well plates in StemMACS PSC-Brew XF medium supplemented with 2.5 µM Y-27632 and StemMACS PSC-Support XF. After 2-4 days, the medium was replaced with StemMACS PSC-Brew XF medium supplemented with 2.5 µM Y-27632. Afterwards, the medium (without Y-27632) was changed every other day. Colonies were visible 7 days after limiting dilution. The first passaging was initiated when colonies covered 40% of the well, or at the latest 10 days after limiting dilution.

For passaging, StemMACS Passaging Solution XF was used. Only a maximum of 10 colonies at a time were passaged, and a separate 96-well plate was used for each round. The cell suspension of a colony was divided into two 96-well plates, one for Sanger sequencing and the other for further expansion. The extraction of gDNA was performed as described above. The genotype was determined in parallel with the expansion. Sanger sequencing results were analysed using EditR (Kluesner *et al*., 2018): Clones with a C-to-T conversion rate of 40-60% were defined as heterozygous, 90-100% as homozygous, and 0-10% as WT. Other populations were considered non-clonal and discarded. The desired clones were expanded stepwise from 48- and 12-well plates to 6-well plates and then cryopreserved.

### Estimation of clones needed to seed

Editing efficiencies were determined by Sanger sequencing and a possible overestimation by up to 10% was considered (Cornean *et al*., 2022; Kluesner *et al*., 2018). The probability for homozygosity (for independent and dependent editing of the alleles and genes, respectively) and the required number of clones to plate were calculated as follows:

● Probability for homozygosity

○ Independent (%) = ((Editing_Gene1_ × Editing_Gene1_) × (Editing_Gene2_ × Editing_Gene2_) × (Editing_Gene3_ × Editing_Gene3_)) × 100
○ Dependent (%) = (Editing_Gene1_ × Editing_Gene2_ × Editing_Gene3_) × 100
● Seeding for x clones = x / (Probability for homozygosity × Recovery) The values are shown in Supplementary Tables S3 and S4.

### Trilineage differentiation

To differentiate hiPSCs into cells of all three germ layers, the StemMACS Trilineage Differentiation Kit (Miltenyi Biotec) was used. hiPSCs were thawed, passaged at 1:4 and 1:6, and then seeded at the recommended cell number in a 12-well plate. For wells intended for immunofluorescence staining, cells were seeded on 18-mm glass dishes. The medium was changed according to the provided protocol and at the same time of day. Cells were harvested for analysis by immunofluorescence and RT-qPCR 7 days after seeding.

### Karyotyping

Genomic DNA was extracted from hiPSC pellets stored at −80°C using the Monarch Genomic DNA Purification Kit (NEB). SNP-array karyotyping was performed as a service by Life & Brain Research Centre (University Hospital of Bonn) using an Illumina BeadArray (≥700K markers). CNVs and regions of homozygosity were assessed from the resulting genotyping data using GenomeStudio (Illumina, V2.0.5) and copy number analysis using cnvPartition (Illumina, v3.2) by the service provider.

### Differentiation of atrial cardiomyocytes

The differentiation and maintenance of atrial CMs was performed as described previously. Atrial cardiomyocyte differentiation was performed in 2D monolayer format, adapted from a previously described protocol (Zhang et al., 2015) with modifications. Briefly, hiPSCs were seeded at 850,000 cells per well in Matrigel-coated 24-well plates in day 0 medium consisting of KO-DMEM supplemented with 1× ITS, 1× Penicillin/Streptomycin, 1× GlutaMAX, 10 ng/ml

FGF2, 10 ng/ml Activin A, 10 ng/ml BMP4, 1 µM CHIR99021, and 10 µM Y-27632. From day 1 onwards, cells were cultured in TS medium (KO-DMEM, 5 µg/ml transferrin, 0.68 ng/ml sodium selenite, 1× Penicillin/Streptomycin, 1× GlutaMAX) supplemented with 250 µM 2-phospho-ascorbate throughout, with the following additional factors: 1 µM CHIR99021 (day 1), 0.2 µM IWP-2 (days 2-3), and 0.5 µM retinoic acid (days 3-4) for atrial specification. From day 5, medium was changed every 2-3 days. On day 15, TS medium was replaced with RPMI supplemented with 1× B27, and cardiomyocytes were purified using the PSC-Derived Cardiomyocyte Isolation Kit (Miltenyi Biotec).

### Immunofluorescence staining

**hiPSC.** For fixation, cells were washed three times with PBS for 5 min each, then fixed with 4% PFA at room temperature. After 10 min of incubation, samples were washed twice with PBS for 5 min each. Fixed cells were stored in PBS + 1% PFA at 4 °C until further processing. Cells were permeabilised with 0.3% Triton in PBS for 30 min at room temperature and then washed three times with PBS for 5 min each. Afterwards, samples were blocked in 1% BSA/0.1% Tween/0.3 M Glycine in PBS for for 60 min at room temperature. The primary antibody was added at the dilutions listed in the key resources table in 1% BSA/0.1% Tween in PBS and incubated overnight at 4°C. The next day, cells were washed three times with PBS for 5 min each. Subsequently, the secondary antibody was incubated in 1% BSA/0.1% Tween in PBS for 60 min at room temperature in the dark. After three washes, DAPI (1.5 µg/ml in PBS) was added, and the cells were incubated for 10 min at room temperature in the dark. Finally, the cells were washed three times and mounted. The samples were dried overnight at room temperature in the dark and stored at 4°C.

**hiPSC-derived cardiomyocytes.** We performed the OCaR2 antibody staining following a previously described protocol (Tsvilovskyy et al., 2024), with some modifications. In brief, cardiomyocytes were fixed as hiPSCs described above, and the fixed cells were permeabilised with ice-cold acetone for 5 min. After washing with PBS for 10 min, 0.1% Tween/PBS was added for 10 min, followed by 50 mM Glycine/PBS for 60 min. For blocking, 1% BSA in PBS was used for 30 min. Afterwards, OCaR2 and cTnT in 1% BSA/PBS were added (key resources table). For OCaR2, antisera were affinity-purified against the C-terminal epitope of OCaR2 (EDSLIENEIHQ) and used at 1:50 dilutions for immunostainings in iPS-derived cardiomyocytes. The plate sealed with parafilm was kept overnight at 4°C. The next day, the samples were washed three times. Next, the secondary antibodies in 1% BSA in PBS were incubated for 60 min at room temperature in the dark. After washing twice with PBS for 10 min each, 1.5 µg/ml DAPI in PBS was added for 10 min. The cells were washed again with PBS (5 min), 1% BSA in PBS (10 min) and PBS (10 min). Finally, the cells were incubated with 10mM Tris/HCl for 10 min and, after drying for 15 min, mounted. The cells were dried overnight at room temperature and then stored at 4°C.

Samples were imaged using a Zeiss Axio Observer Z1 microscope (Zeiss) with ZEN Pro software (Zeiss) and a 63x/1.4 NA (oil) objective. The images were further edited with the Fiji distribution of ImageJ (version 2.16.0/1.54p) (Schindelin et al., 2012). Antibodies and their dilution are listed in the key resources table.

### Analysis of OCaR2 signal (Immunofluorescence staining)

The differentiation of hiPSCs into atrial CMs was carried out in three separate wells for both WT and dKO. The aCMs were passaged onto separate 18-mm glass coverslips and stained. From each coverslip, a microscopic image was taken and further analysed identically with Fiji. In Fiji, for each image, the colour thresholds for cTnT and OCaR2 were set identically, and the cTnT ROI was transferred to the OCaR2 image. Then, the percentage of OCaR2 area that overlapped with the cTnT-ROI was measured.

### Reverse-transcription quantitative PCR (RT-qPCR)

Total RNA was extracted using the RNeasy Mini Kit (Qiagen) with on-column DNase digestion (RNase-Free DNase Set, Qiagen). For cDNA synthesis, 1 µg of RNA was reverse-transcribed using the SensiFAST cDNA Synthesis Kit (Meridian Bioscience) following the manufacturer’s protocol. qPCR reactions were assembled using the SensiFAST SYBR Hi-ROX Kit (Meridian Bioscience) using 50 ng cDNA as template per reaction and run on a LightCycler 96 instrument (Roche) with the following cycling conditions: 95°C for 10 min, followed by 40 cycles of 95°C for 10 s, 60°C for 10 s, and 72°C for 30 s, with a subsequent melting curve analysis (95°C for 10 s, 65°C for 60 s, 97°C for 1 s). All samples were measured in technical duplicate. Primer efficiencies were determined from standard curves using three-point serial dilutions of cDNA (50, 10, and 2 ng input). Relative expression was calculated using the efficiency-corrected method (Pfaffl, 2001), normalised to the arithmetic mean of *GAPDH* and *ACTB* reference gene expression. Primer sequences are listed in Supplementary Table S5. For pluripotency assessment (*SOX2, OCT4, NANOG*, and trilineage markers), clones were considered pluripotent when expression levels exceeded 50% of wild-type levels.

### Mycoplasma testing

Before transfections or cell banking, all hiPSC KO clones were tested for mycoplasma as previously described (Uphoff and Drexler, 2004) using Q5 Hot Start Polymerase and a previously described internal control (Uphoff and Drexler, 2002).

### Targeted amplicon sequencing and data analysis

Samples were prepared for targeted amplicon sequencing either by two-step PCR with multiplexed indexing followed by sequencing (GeneWiz/Azenta Life Sciences), as previously described (Ong *et al*., 2026), or by Amplicon-EZ (GeneWiz/Azenta Life Sciences), as previously described (Cornean *et al*., 2022). For multiplexed samples, locus-specific primers carried partial Illumina adapter overhangs (listed in Supplementary Table S5), and amplicons from different target loci within each biological replicate were pooled into a single submission and demultiplexed bioinformatically by mapping to the respective reference sequences. NGS data (in FASTQ format) were processed using CRISPResso2 v2.2.12 (Clement et al., 2019) in batch mode with base editing parameters (--min_average_read_quality 30, --quantification_window_size 20, --quantification_window_center −10, --base_editor_output). C-to-T editing frequencies across the protospacer were extracted from the Nucleotide_percentage_summary output file. Indel frequencies were extracted from the quantification_of_editing_frequency output file and calculated as previously described (Cornean *et al*., 2022). Downstream analysis and visualisation were performed in R (v4.3.1).

### Whole-genome sequencing sample preparation and sequencing

Genomic DNA was extracted from hiPSC pellets (wild-type parental line, O2-sKO, and O2T2-dKO clones) stored at −80 °C using the Monarch Genomic DNA Purification Kit (NEB). DNA was quantified using the Qubit dsDNA BR Assay. DNA integrity was confirmed by Agilent Genomic DNA ScreenTape analysis. Whole-genome sequencing was performed by Azenta Life Sciences (GENEWIZ Germany) using their Short-Read Human WGS service on an Illumina NovaSeq platform (2 × 150 bp paired-end reads, 30x target coverage).

### Whole-genome sequencing: bioinformatics analysis

Reads were aligned to the GRCh38 reference genome (Ensembl primary assembly) using BWA-MEM and processed through the nf-core/sarek pipeline (v3.7.1). Duplicate reads were marked using GATK MarkDuplicates, and base quality score recalibration (BQSR) was performed using GATK BaseRecalibrator. Analysis was restricted to canonical chromosomes (chr1-22, X, Y). Somatic variant calling was performed using GATK Mutect2 in tumour-normal mode, with the parental wild-type hiPSC line as the matched normal, gnomAD allele frequencies for population filtering, and a 1000 Genomes panel of normals. Mutect2 filtering included cross-sample contamination estimation (CalculateContamination) and orientation bias correction (LearnReadOrientationModel). Structural variants were called using Manta, and CNVs using CNVKit. This filtering approach for somatic variants was adapted from earlier work (Kim *et al*., 2025) for comprehensive off-target analysis of base editors using whole-genome sequencing: PASS filter status, biallelic variants only, tumour read depth ≥15, tumour alternative allele reads ≥5, and normal alternative allele reads = 0.

### Off-target site prediction and variant classification

Potential guide RNA off-target sites were predicted using Cas-OFFinder v2.4 (Bae *et al*., 2014) with the following parameters: ≤3 mismatches, ≤1 DNA bulge, no RNA bulge, against the GRCh38 reference genome. Guide sequences queried were CATAGTCCCAGGCCACCTTC (g1/O2-3, targeting *OCaR2*/*TMEM63B* at chr6:44,135,019-44,135,042) and AACCCTGCAAGATAGACGGG (g3/T2-1, targeting *TPC2*/*TPCN2* at chr11:69,057,535-69,057,558), both with NGG PAM. This yielded 262 predicted sites for g1 and 158 for g3 (420 total, 212 after merging overlapping windows).

Observed somatic variants were classified into four tiers based on their overlap with predicted off-target sites and consistency with cytosine base editor activity. Tier 1: C>T or G>A variants at a predicted off-target site, falling within the evoBE4max editing window (protospacer positions C3-C10). Tier 2: variants at a predicted off-target site but outside the canonical editing window. Tier 3: C>T or G>A variants genome-wide that do not coincide with any predicted off-target site (stochastic CBE background). Tier 4: non-CBE substitutions not at predicted sites (non-editor background). Tier 1, Tier 2, and select Tier 3 variants were visually inspected using the Integrative Genomics Viewer (IGV) to confirm read-level support, including assessment of mapping quality, read strand balance, and absence of artefact signatures.

An orthogonal, prediction-free guide homology analysis was conducted as sensitivity analysis: for all HIGH-impact variants (VEP consequence), ±25 bp flanking sequence was extracted and a 20-mer sliding window was aligned against g1 and g3 guide sequences to determine the minimum mismatch count independent of Cas-OFFinder predictions.

### Variant annotation and mutational signature analysis

Variants were annotated with Ensembl VEP v115 using ClinVar (2025 release) and COSMIC v103 databases. Functional impact was assessed using VEP consequence and IMPACT fields. For trinucleotide context analysis, the 5’ and 3’ flanking bases of each C>T or G>A SNV were extracted from the GRCh38 reference genome. G>A variants were reverse-complemented to report all variants on the C>T pyrimidine strand. Counts were computed across 16 possible 5’-N[C>T]N-3’ trinucleotide contexts.

C-to-T enrichment was quantified as an odds ratio relative to a null model assuming equal probability (1/6 ≈ 16.7%) for each of six pyrimidine-normalised substitution classes (C>A, C>G, C>T, T>A, T>C, T>G). 95% confidence intervals were computed using the log method. Variant overlap between clones was determined by exact position match (chromosome:position:reference:alternative allele). Allele fraction clonality was assessed by classifying variants with AF ≥ 0.4 as clonal (present at or near the time of editing) and AF <0.4 as subclonal (arising during subsequent expansion).

Circular genome plots were generated using the R package circlize, integrating chromosome ideograms, predicted off-target windows, observed Tier 1+2 variants, CNV segments from CNVKit, and structural variant breakpoint links from Manta. Variant landscape plots, mutation spectra, filtering funnels, and statistical analyses were generated using R with ggplot2.

### Software versions in genome analysis

All analyses were performed using the following software supported by Python (v3.9.6) preprocessing script: nf-core/sarek v3.7.1 (Ewels et al., 2020; Garcia et al., 2020; Hanssen et al., 2024), incorporating GATK Mutect2 (Benjamin et al., 2019), BWA-MEM (Li, 2013), Manta (Chen et al., 2016), and CNVKit (Talevich et al., 2016). Variant annotation was performed with Ensembl VEP v115 (McLaren et al., 2016) using ClinVar (Landrum et al., 2025) and COSMIC v103 (Tate et al., 2019). Off-target sites were predicted with Cas-OFFinder v2.4 (Bae *et al*., 2014). Circular genome plots were generated using the R package circlize (Gu et al., 2014). Variant landscape plots, mutation spectra, filtering funnels, and statistical analyses were generated using R with ggplot2 (v3.5.1) (Wickham, 2016). WGS variant inspection was conducted manually using the IGV-Web application (v2.4.3) (Robinson et al., 2011).

### Statistics and graphics

All data in the text are displayed as mean value (± standard deviation (SD)). Data were visualised in R (Version 4.3.1) using the IDE RStudio (Version 2023.06.1+524) or Visual Studio Code (Version 1.113.0) with the packages listed in the key resources table. Figures were created in Affinity Designer 2 and Adobe Illustrator, respectively, with some icons imported from BioRender.

### Ethics declaration

The study was approved by the Ethics Committee of the Heidelberg Medical Faculty (case numbers: S-224/2022 (approved 28/04/2022) and S-148/2022 (approved 17/03/2022)).

## RESOURCE AVAILABILITY

### Lead contact

Requests for further information and resources should be directed to and will be fulfilled by the lead contact, Alex Cornean.

### Materials availability

Plasmids pEF1α_evoBE4max (Addgene #255768) and pEF1α_CBE-T1.46 (Addgene #255777) have been deposited at Addgene and will be available upon publication. Any other materials and reagents will be made available upon reasonable request.

### Data and code availability

Oligonucleotide and sgRNA sequences are listed in the key resources table and Supplementary Table S5. Targeted amplicon sequencing data have been deposited at the NCBI Sequence Read Archive under BioProject PRJNA1458718. Sanger sequencing traces, SNP-array karyotyping data, and original analysis code have been deposited at Zenodo (https://doi.org/10.5281/zenodo.19652563). The WGS off-target and mutational burden analysis pipeline has been deposited at Zenodo (https://doi.org/10.5281/zenodo.20506479) and is also publicly available on GitHub (https://github.com/Alex-Cornean/wgs-offtarget-burden-CBE). Processed somatic variant calls from whole-genome sequencing (VCF format, germline variants excluded, normal sample data removed) will be deposited at Zenodo upon publication. Raw WGS sequencing reads are not publicly available owing to donor consent restrictions under the cell line provider’s material transfer agreement. Any additional information required to reanalyse the data reported in this paper is available from the lead contact upon reasonable request.

## Supporting information

contains Figures S1-S8, Tables S1-S4

## ACKNOWLEDGMENTS

Sayari Bhunia is a member of HBIGS, the Heidelberg Biosciences International Graduate School. We thank the entire Freichel lab for their constructive feedback on the manuscript. We thank Kristin Rädecke and Amelie Obermeier for providing differentiated atrial cardiomyocytes and Tinatini Tavhelidse-Suck for her support in coordinating the karyotyping. We are grateful to Veit Flockerzi for his support in producing the anti-OCaR2 antibody. We thank the Flow Cytometry & FACS Core Facility (FFCF) at ZMBH, Heidelberg University, for cell sorting support. This research was funded by the German Research Foundation (DFG) through the Collaborative Research Centres CRC1550 (FKZ 464424253 project B09 to MF), and CRC1328 (FKZ 335447717, project A21 to MF), as well as the FOR5705 (523862973, FR 1638/6-1, Neuroflame project P03 to MF), and the DZHK (German Centre for Cardiovascular Research), Heidelberg/Mannheim and the BMBF (German Ministry of Education and Research). The authors further acknowledge support by the state of Baden-Württemberg through bwHPC and the DFG through grant INST 35/1597-1 FUGG. Vanessa Kirschner was supported by a MD Fellowship from the Collaborative Research Centre CRC1550. During the preparation of this work, the authors used AI-assisted technologies (Grammarly and Claude by Anthropic) to improve readability and language. After using these tools, the authors reviewed and edited the content as needed and take full responsibility for the publication’s content.

## AUTHOR CONTRIBUTIONS

V.K.: Conceptualisation, Data curation, Formal analysis, Investigation, Methodology, Validation, Visualisation, Writing – original draft, Writing – review & editing.

J.H.: Investigation, Data curation, Validation, Writing – review & editing. C.R.: Investigation, Validation, Writing – review & editing.

S.B.: Investigation, Validation, Writing – review & editing. R.K.: Resources, Writing – review & editing.

T.S.: Resources, Writing – review & editing.

M.F.: Conceptualisation, Funding acquisition, Project administration, Resources, Writing – review & editing.

A.C.: Conceptualisation, Data curation, Formal analysis, Software, Methodology, Project administration, Supervision, Visualisation, Writing – original draft, Writing – review & editing. *All authors read and approved the final manuscript*.

## DECLARATION OF INTERESTS

The authors declare no competing interests.

## SUPPLEMENTAL INFORMATION

**Document S1**. Figures S1-S8, Tables S1-S4

**Table S5**. sgRNA cloning oligonucleotides, amplicon sequences, sgRNA sequences, Gibson assembly oligonucleotides, target amplicon PCR oligonucleotides, and RT-qPCR primers, related to Experimental Procedures

## Notes

### Competing Interest Statement

The authors have declared no competing interest.

