## Supplementary material for "Cytosine base editing workflow for quality-controlled multiplex-knockout hiPSC lines": contains Figures S1-S8, Tables S1-S4

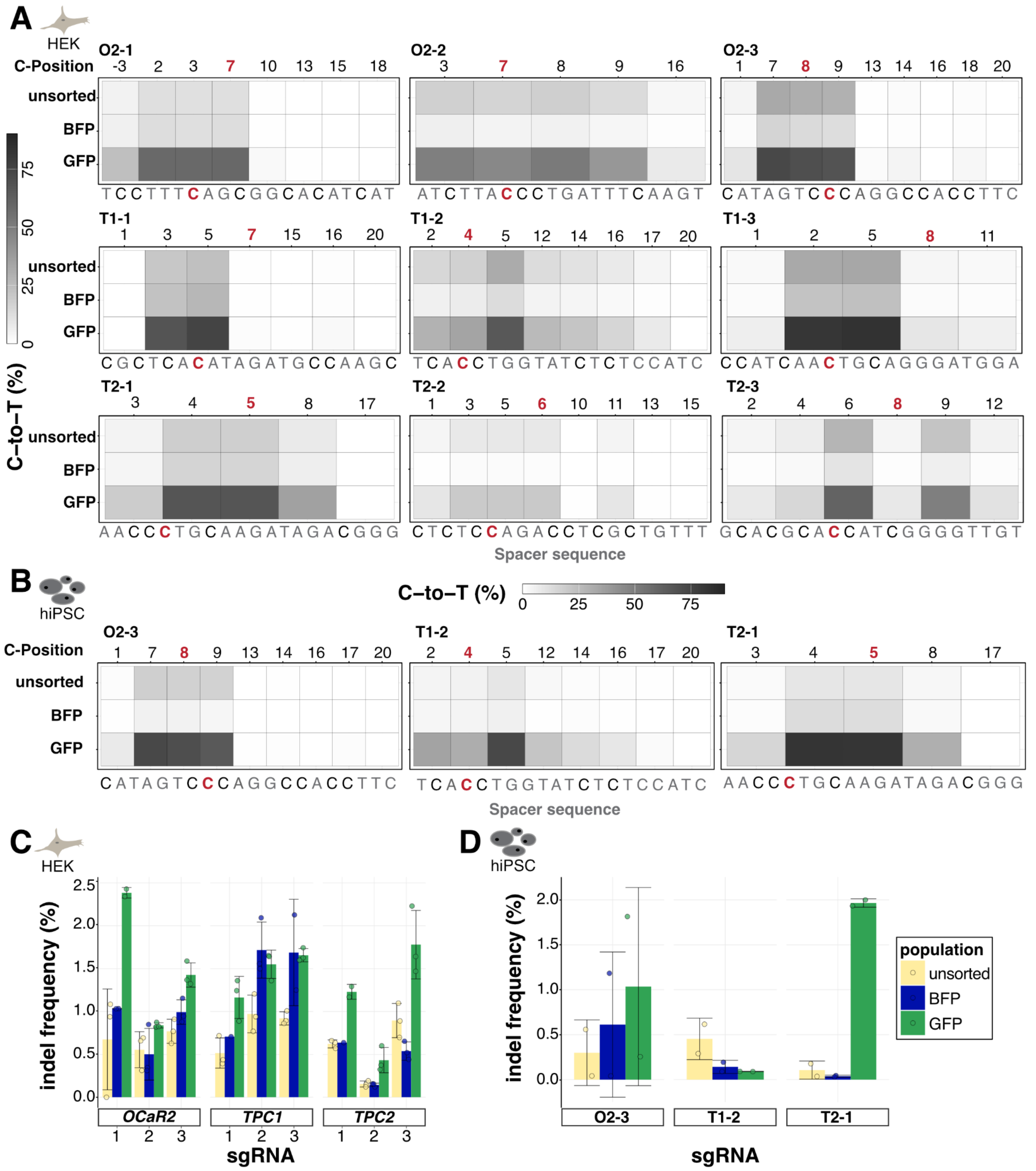


**Figure S1 Heatmaps of C-to-T editing and barplots of indel frequency in HEK293T cells and hiPSCs, related to Figures 2 and 3:**
**(A)** Percentage of C-to-T editing of every cytosine along the protospacer for all candidate sgRNAs for unsorted, BFP and GFP-positive HEK293T cells. The number above indicates the position of each cytosine.
**(B)** Heatmaps of C-to-T editing in all three populations for the selected sgRNAs in hiPSCs. The respective spacer sequence is written below each heatmap. The target cytosine is marked in red.
**(C-D)** Percentage of indels of all three populations in HEK293T cells for all sgRNA candidates **(C)** and in hiPSCs for the selected sgRNAs **(D).** Data were obtained by amplicon deep sequencing. Heatmaps (A-B) show the mean of independent replicates; barplots (C-D) show mean ± SD. For A and C: n=3 per population, except: n=2 for O2-1 BFP and GFP, T1-3 BFP, T2-1 GFP; n=1 for T1-1 BFP, T2-1 BFP; for B and D: n=2 per population.

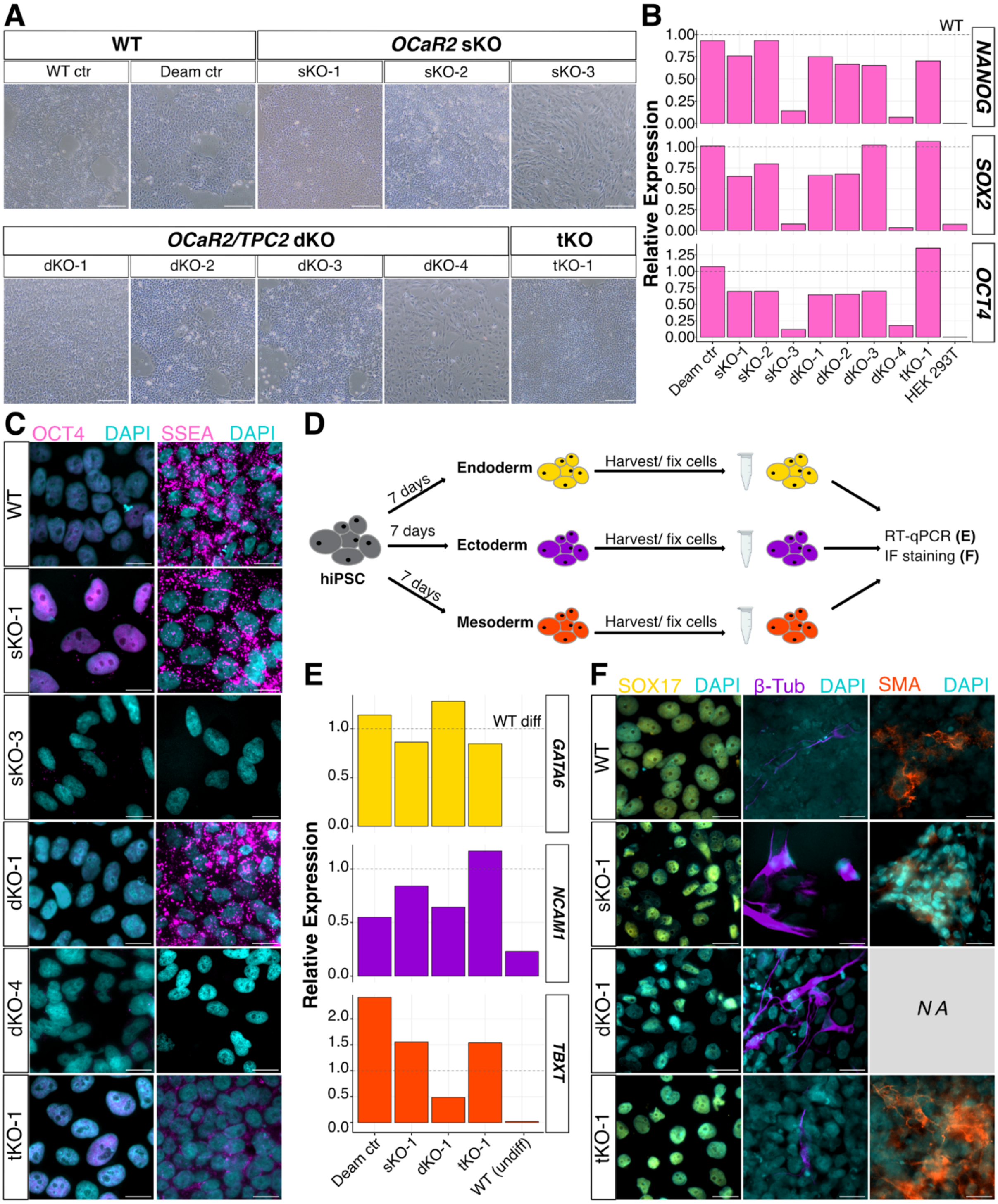


**Figure S2: Characterisation of hiPSC KO clones, related to Figures 4 and 5.**
**(A)** Phase contrast microscopy images of all homozygous hiPSC KO clones and wild-type controls. The images were taken with an EVOS XL Core microscope; scale bar = 200 µm.
**(B)** RT-qPCR results of pluripotency markers (*NANOG*, *SOX2*, *OCT4*) of all homozygous hiPSC KO clones (single measurement). Relative expression was normalised to wild-type hiPSCs (dotted line). HEK293T cells were used as negative control.
**(C)** Immunofluorescence staining of representative samples for *OCaR2* sKO, *OCaR2/TPC2* dKO, *OCaR2/TPC1/TPC2* tKO, and wild-type hiPSCs. The overlay of DAPI (cyan) and OCT4 or SSEA (magenta) is shown; scale bar = 20 µm.

**Figure S2 (continued):**
**(D)** Schematic procedure of trilineage differentiation: Wild-type, Deaminase control, and one clone each of *OCaR2* sKO and *OCaR2/TPC2* dKO and *OCaR2/TPC1/TPC2* tKO were differentiated into early stages of all three germ layers, Ectoderm, Endoderm and Mesoderm. After 7 days, the cells were harvested for RT-qPCR analysis or fixed for immunofluorescence staining.
**(E)** RT-qPCR results of Trilineage Differentiation: Relative expression was normalised to differentiated wild-type cells (dotted line). The normalised expression of *GATA6* (Endoderm), *TBXT* (Mesoderm) and *NCAM1* (Ectoderm) is shown. n=2 (biological replicates).
**(F)** Immunofluorescence staining of markers for Trilineage Differentiation: depicted are representative samples of the overlay of SOX17 (Endoderm, yellow), β-Tubulin III (Ectoderm, purple) or SMA (Mesoderm, orange) with DAPI (cyan); scale bar = 20 µm. ctr, control; WT, wild-type; IF, immunofluorescence; Deam ctr, Deaminase control.

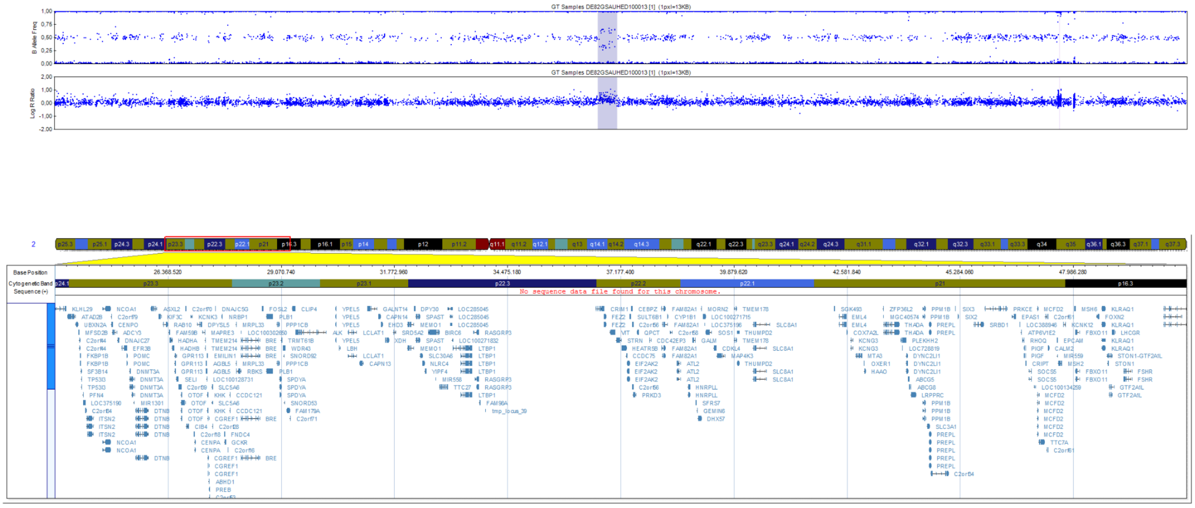

**Figure S3: SNP-array karyotyping of clone *OCaR2* sKO-1 showing a 475 kbp duplication on chromosome 2p, related to Figures 4 and 6.** Upper panel: B-allele frequency; middle panel: log R ratio - the duplication region is highlighted; lower panel: chromosome 2 overview with zoom in on genes encoded in affected regions;

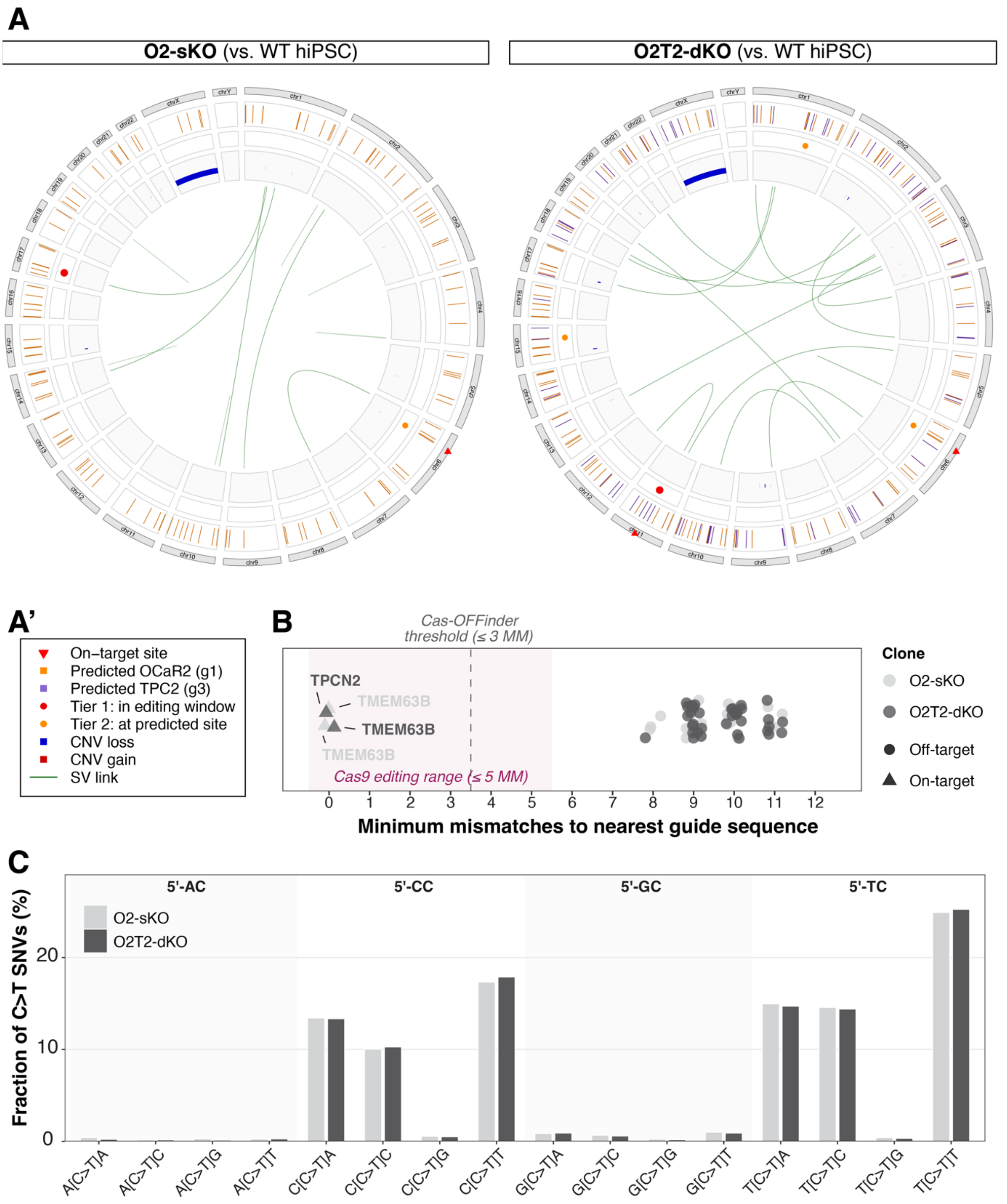


**Figure S4: Genome-wide variant distribution, guide homology analysis, and trinucleotide context, related to Figure 6.**
**(A)** Genome-wide circos plots of somatic variants in the O2-sKO (left) and O2T2-dKO (right) clones called against the parental wild-type hiPSC line. Outer tracks: chromosome ideograms with on-target sites (red triangles). Middle tracks: Cas-OFFinder-predicted off-target windows for *OCaR2*/g1 (orange squares) and *TPC2*/g3 (purple squares). Inner symbols: Tier 1 variants (red circles), Tier 2 variants (orange circles), CNV losses (blue bars), CNV gains (red bars), and structural variant links (green arcs). No structural variants or CNVs overlap predicted off-target sites.
**(A')** Legend for circos plot symbols and annotation tracks.

**Figure S4 (continued):**
**(B)** Orthogonal guide homology analysis. For all 66 HIGH-impact variants across both clones, the minimum number of mismatches to the nearest guide sequence (g1 or g3) was determined by sliding-window alignment. Variants at on-target loci (*TMEM63B*, *TPCN2*) show 0 mismatches (labelled). All other HIGH-impact variants have ≥8 mismatches, far beyond the established range of Cas9-mediated editing (shaded region, 0-5 mismatches; (Hsu et al., 2013; Tsai et al., 2015)), confirming that no HIGH-impact variant outside on-target loci is guide-directed.
**(C)** Trinucleotide context of C>T/G>A variants (16-bin, 5'-N[C>T]N-3'). TC and CC motifs account for ca. 40% and ca. 41-42% of C>T variants, respectively, consistent with the rAPOBEC1 deaminase in evoBE4max. AC contexts are depleted (ca. 1%), mirroring the disfavoured AC dinucleotide observed at on-target sites during editor benchmarking (**Figure 1A**). GC contexts account for <3%.

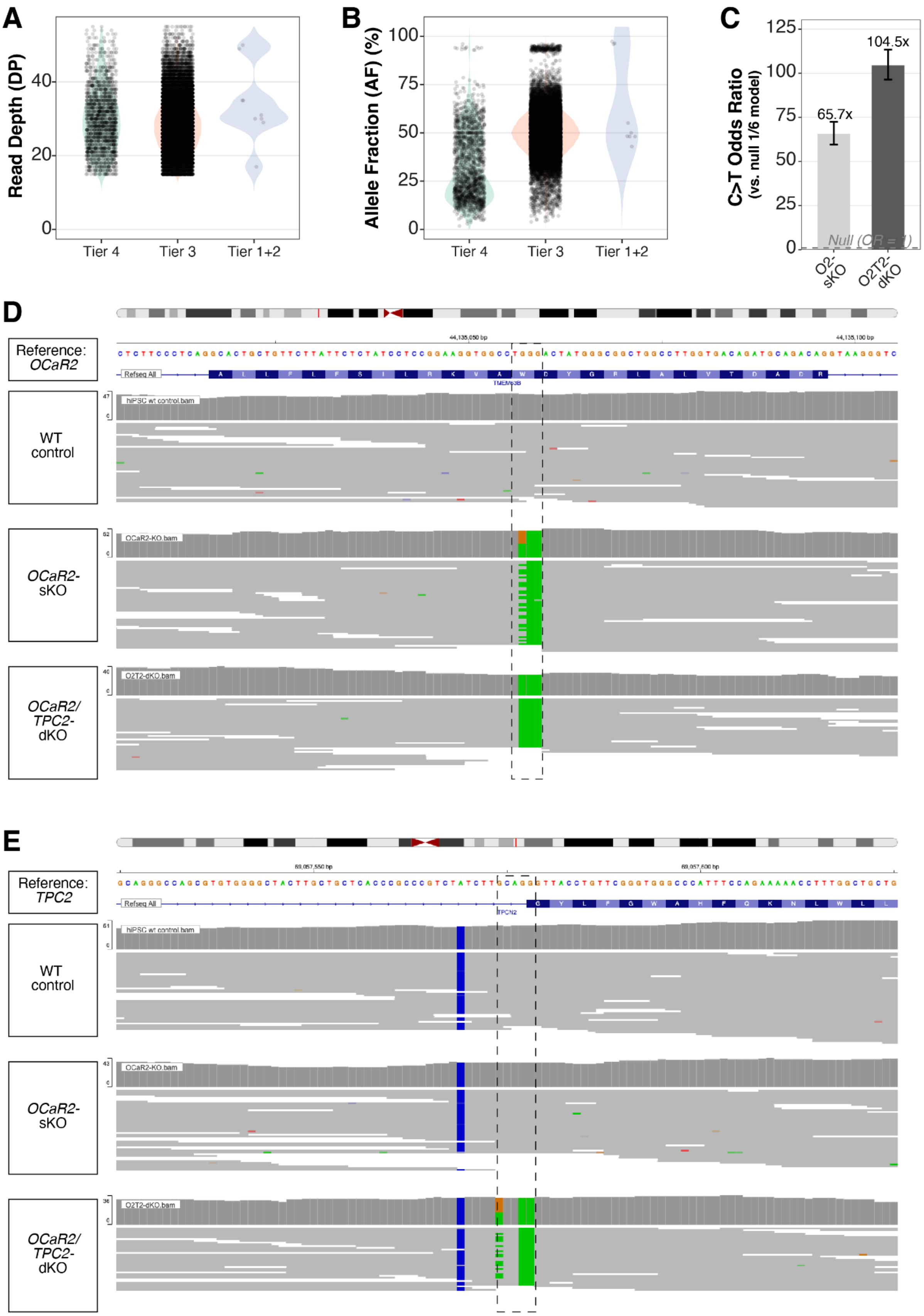


**Figure S5: Read depth, allele fraction distributions, and C>T enrichment, related to Figure 6.**

**Figure S5 (continued):**
**(A)** Violin plots with overlaid data points showing read depth (DP, left) and allele fraction (AF, right) stratified by variant tier. Tier 1+2 on-target variants show near-homozygous AF (>0.95); the single genuine off-target Tier 1 variant is heterozygous (AF 0.48). Approximately 84% of Tier 3 background variants are clonal (AF ≥ 0.4), indicating CBE-mediated deamination at or near the time of editing.
**(B)** C>T odds ratio with 95% confidence intervals for both clones (O2-sKO: 65.7, 95% CI 59.6-72.5; O2T2-dKO: 104.5, 95% CI 96.4-113.4), computed relative to a null model of equal probability (1/6) for each of 6 pyrimidine-normalised substitution classes. Dashed line: null expectation (OR = 1). IGV screenshots of *OCaR2* (**C**) and *TPC2* (**D**) on-target sites in WGS data.

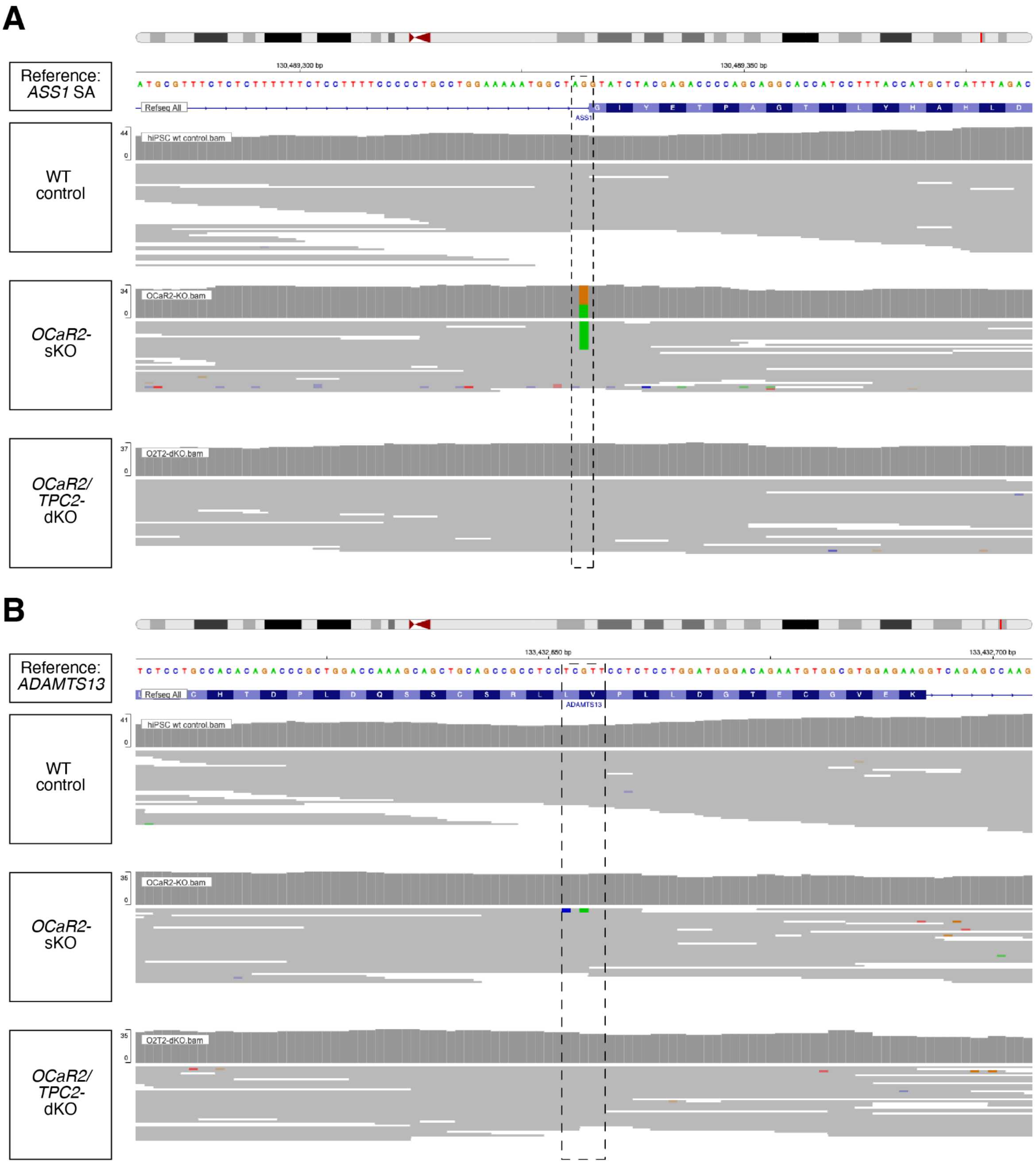


**Figure S6: IGV visualisation of ClinVar-flagged variants, related to Figure 6.**
**(A)** IGV screenshot of the *ASS1* splice-acceptor variant (chr9:130,489,332, G>A) in the O2-sKO clone. This ClinVar likely-pathogenic variant is heterozygous (AF = 0.50), classified as Tier 3, and has 9 mismatches to the nearest guide, a stochastic CBE-mediated deamination event.
**(B)** IGV screenshot of the *ADAMTS13* missense variant (chr9:133,432,654, G>A) in the O2-sKO clone (AF = 0.08, ClinVar conflicting interpretations).

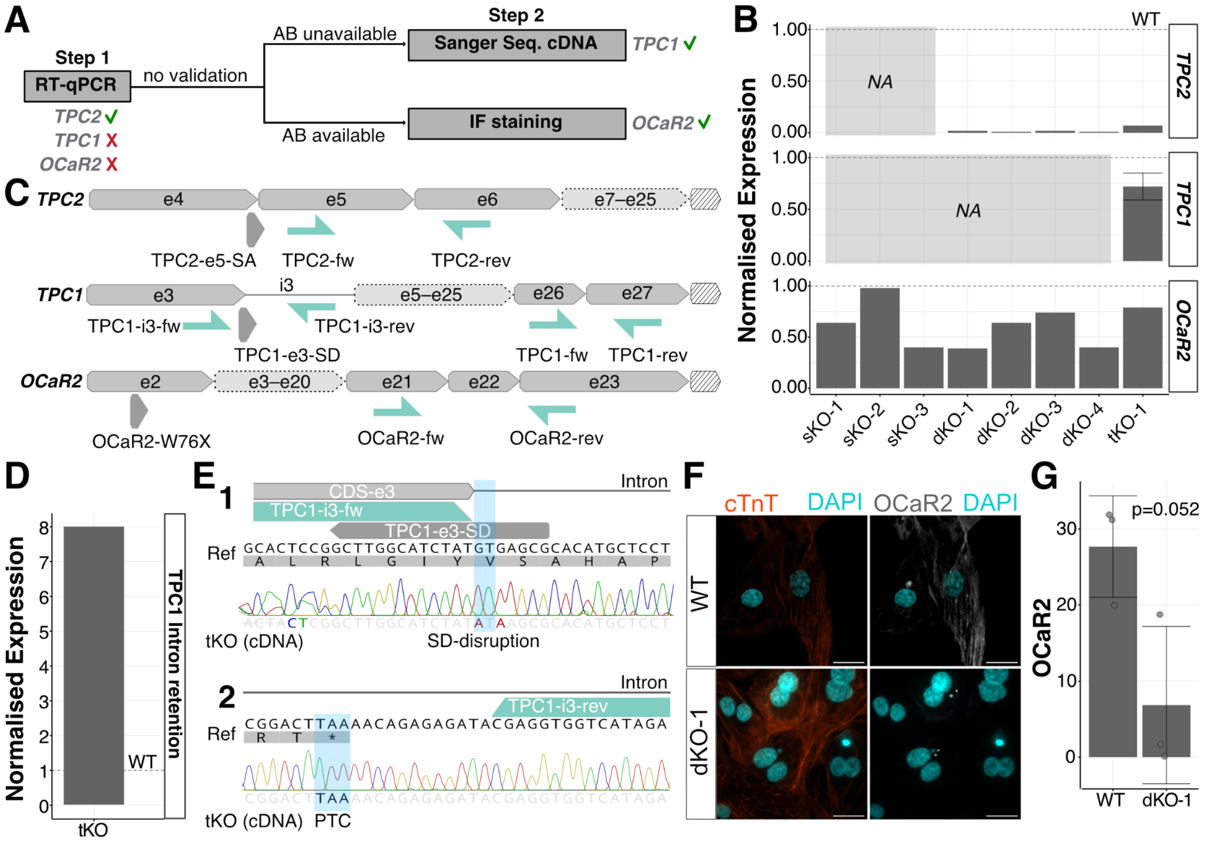


**Figure S7: Transcript- and protein-level analyses for the validation of predicted loss-of-function alleles, related to Figures 4 and 5.**

**(A)** Validation process of each gene: RT-qPCR was performed for all three genes. If the result was unclear, either immunofluorescence staining (*OCaR2*) or Sanger sequencing for cDNA (*TPC1*) followed, depending on antibody availability for the target protein.

**(B)** RT-qPCR results for the expression of *OCaR2, TPC1* and *TPC2* in all homozygous KO clones (single measurement). The relative expression was normalised to wild-type hiPSCs (dotted line).

**(C)** Overview of sgRNAs (dark grey) and RT-qPCR primer pairs (light green). The hatched arrow indicates the 3'UTR.

**(D)** *TPC1* intron retention levels of the tKO normalised to wild-type hiPSCs (dotted line) were quantified using exon 3-intron 3 spanning primers.

**(E)** Sanger sequencing results of tKO cDNA for the SD-disruption (1) and the following PTC (2): the coding sequence of exon 3 and the following intron are indicated in dark grey, the primers used in light green and the sgRNA used for the KO is shown below. Nucleotides that deviate from the reference are marked in colour. The resulting SD-disruption and following stop codon are highlighted in blue.

**(F)** Immunofluorescence staining of OCaR2: for the identification of cardiomyocytes, cTnT (orange) was used. OCaR2 is depicted in gray. The overlay with DAPI (cyan) is shown.

**(G)** Analysis of immunofluorescence staining: Percentage of area with OCaR2 signal normalised to cTnT signal. Each data point corresponds to one ROI measured from a separate staining sample (one ROI per coverslip). A Student's (independent) *t*-test was performed against WT ($p=0.052$). Values are shown as mean ± SD. AB, antibody; SD, splice donor; PTC, premature termination codon.

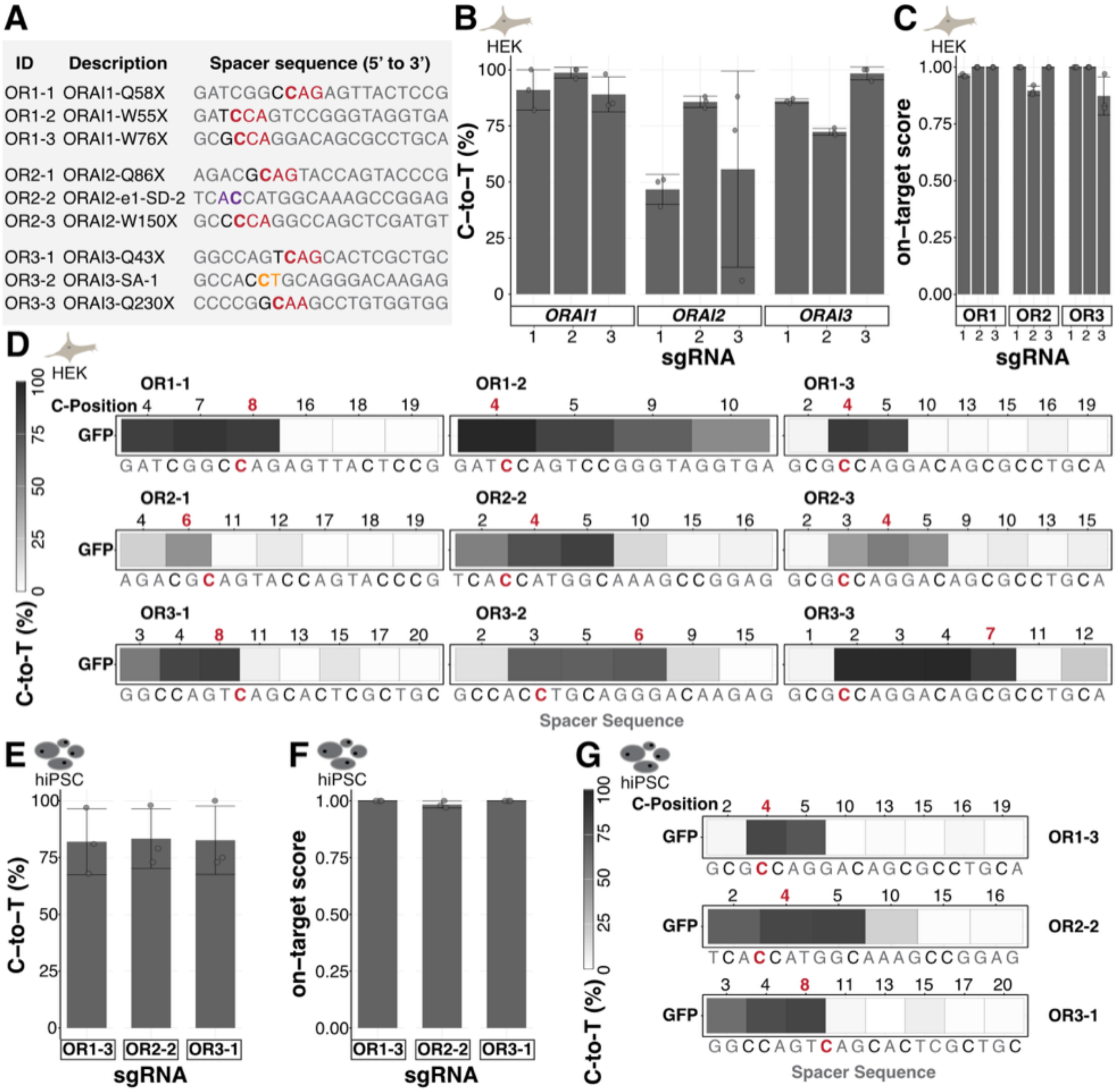


**Figure S8: Application of the workflow to *ORAI* genes, related to Figure 7.**
**(A)** Design of candidate sgRNA to create *ORAI1, ORAI2* and *ORAI3* KO cells via nonsense (X, dark red), splice donor (SD, purple), or splice acceptor (SA, orange) mutation facilitated by CBE. The affected codon or splice site within the protospacer is marked with the corresponding colour, the target cytosine in bold, the cytosine context in the shape of the preceding nucleotide in black, and the remaining protospacer sequence in grey.
**(B)** Peak editing after FACS in the GFP-positive population of HEK293T cells for all sgRNA candidates.
**(C)** Ratio of editing at the target cytosine and peak editing for GFP-positive HEK293T cells. **(D)** Percentage of C-to-T editing at all cytosines in the protospacer for HEK293T cells. The numbers indicate the position of each cytosine within the protospacer. The target cytosine is marked in red. The sequence of the protospacer for each sgRNA is written below.
**(E-F)** Peak editing **(E)** and on-target score **(F)** of the GFP-positive population of selected sgRNAs in hiPSCs are shown.
**(G)** Heatmap of C-to-T editing at every cytosine within the protospacer for the selected sgRNAs in hiPSCs. Each experiment was performed with three independent replicates (n=3). Values are shown as mean ± SD and the data obtained by Sanger sequencing.

**Table S1: Details on SNP-array Karyotyping, related to Figure 4 and 5**

| **Sample** | **Result** | **Details** |
|---|---|---|
| Deaminase ctr | No copy number events detected | |
| sKO-1 | Duplication | 475 kbp on chr 2p |
| sKO-2 | No copy number events detected | |
| dKO-1 | No copy number events detected | |
| dKO-2 | No copy number events detected | |
| dKO-3 | Copy number gain involving a hotspot region | 1051 kbp on chr9 |
| tKO-1 | No copy number events detected | |

**Table S2**: Transfection conditions: Plasmid amounts per well of a 6-well plate*. Related to methods

| **Condition** | **CBE (ng)** | **sgRNA (ng each)** | **BFP (ng)** | **Lipofectamine Stem (µl)** | **Total DNA (ng)** |
|---|---|---|---|---|---|
| sKO | 2,700 | 900 (1x) | 900 | 12 | 4,500 |
| dKO | 2,700 | 900 (2x) | 900 | 12 | 5,400 |
| tKO (evoBE4max) | 2,700 | 900 (3x) | 900 | 12 | 6,300 |
| tKO (CBE-T1.46) | 2,100 | 300 (3x) | 300 | 10 | 3,300 |

*In all conditions, Opti-MEM was added to a total reaction volume of 330 µl.

**Table S3:** Calculation of the required cell seeding based on cell recovery and editing efficiencies. Related to Methods

| KO type | Allele editing mode | Recovery | Locus | GFP-pool editing efficiency | Corrected efficiency | Probability for homozygosity | Required seeding for x clones |
|---|---|---|---|---|---|---|---|
| sKO | Ind. | 7.5% | *OCaR2* | 35.0% | 25.0-35.0% | 6.3-12.3% | 326-635 |
| x=3 | Dep. | 7.5% | *OCaR2* | 35.0% | 25.0-35.0% | 25.0%-35.0% | 115-160 |
| dKO | Ind. | 7.5% | *OCaR2*<br>*TPC2* | 40.0%<br>69.0% | 30.0-40.0%<br>59.0-69.0% | 3.1-7.6% | 702-1721 |
| x=4 | Dep. | 7.5% | *OCaR2*<br>*TPC2* | 40.0%<br>69.0% | 30.0-40.0%<br>59.0-69.0% | 17.7-27.6% | 194-302 |
| tKO | Ind. | 7.5% | *OCaR2*<br>*TPC1*<br>*TPC2* | 48.0%<br>40.0%<br>71.0% | 38.0-48.0%<br>30.0-40.0%<br>61.0-71.0% | 0.5-1.9% | 702-2667 |
| x=1 | Dep. | 7.5% | *OCaR2*<br>*TPC1*<br>*TPC2* | 48.0%<br>40.0%<br>71.0% | 38.0-48.0%<br>30.0-40.0%<br>61.0-71.0% | 6.0-12.1% | 111-223 |
| tKO*<br>x=1 | Ind. | 7.5% | *OCaR2*<br>*TPC1*<br>*TPC2* | 17.0%<br>19.0%<br>57.0% | 7.0-17.0%<br>9.0-19.0%<br>47.0-57.0% | 0.3-1.8% | 725-4503 |

Ind. = Independent; Dep. = Dependent.

x: Indicates the actually isolated number of homozygous clones.

* This initial attempt for tKO generation was discontinued.

** Recovery rate (7.5%) was estimated from the observed yield of single-cell clones after limiting dilution across sKO and dKO experiments. Editing efficiencies were derived from Sanger sequencing and corrected by subtracting 10% to account for potential overestimation by EditR-based sequence deconvolution (Kluesner et al. 2018; Cornean et al. 2022). Two allele-editing models were considered: an independent model, in which homozygosity probability equals the product of per-locus homozygosity probabilities, and a dependent model, in which alleles within each locus are edited co-dependently. The dependent model was favoured because observed clone genotypes were predominantly wild-type or homozygous, with few heterozygous clones, and because back-calculated editing efficiencies from actual clone yields (42.9% for sKO, 26.7% for dKO) were more consistent with dependent than independent allelic editing. Under these assumptions, the initial tKO attempt (tKO*) would have required seeding 725-4,503 wells to obtain a single triple-homozygous clone.

**Table S4:** Statistics of the generated KO hiPSC lines. Probability for homozygosity was calculated as the ratio of homozygous KO clones and recovered clones (minus the discarded populations). Related to Methods.

| | Seeded clones | Recovered (discarded) | Recovery | Homozygous KO clones | Probability for homozygosity |
|---|---|---|---|---|---|
| *OCaR2* sKO | 96 | 8 (1) | 7.3% | 3 | 42.9% |
| *OCaR2/TPC2* dKO | 288 | 21 (6) | 5.2% | 4 | 26.7% |
| *OCaR2/TPC1/TPC2* tKO | 192 | 29 (10) | 9.9% | 1 | 5.3% |

**Table S5, separate Excel sheet:** for sgRNA cloning oligonucleotides, amplicon sequences, sgRNA sequences, Gibson assembly oligonucleotides, target amplicon PCR oligonucleotides, and RT-qPCR primers
